# Spinal nociceptive denervation impedes subsequent chronic autonomic remodeling after myocardial infarction in male swine

**DOI:** 10.1101/2025.03.28.645120

**Authors:** Valerie Y.H. van Weperen, Jonathan D. Hoang, Neil Jani, Shail Avasthi, Christopher A. Chan, Kuan Cao, Zulfiqar A. Lokhandwala, Maryam Emamimeybodi, Karim Atmani, Marmar Vaseghi

## Abstract

After chronic myocardial infarction (MI), pathological autonomic remodeling, including vagal dysfunction and sympathoexcitation, predisposes to ventricular arrhythmias (VT/VF). However, what underlies this functional and structural remodeling remains unknown. We hypothesized that sympathetic nociceptive afferent signaling initiates and perpetuates these pathological autonomic changes. We employ cervicothoracic epidural resiniferatoxin (RTX) to ablate spinal nociceptive neurons in male pigs before MI, and assessed autonomic and electrophysiological function four-to-six weeks post-infarction. Compared to vehicle-treated infarcted animals, epidural RTX attenuates the loss of vagal tone and baroreflex sensitivity, reduces spinal cord inflammation, glial activation, and circulating stress and inflammatory markers, and stabilizes electrophysiological parameters, lowering VT/VF inducibility. In a separate cohort, acute C7–T1 nociceptive afferent ablation after chronic MI acutely restores vagal function and decreases VT/VF inducibility. This study demonstrates that cervicothoracic spinal nociceptive afferents significantly contribute to MI-induced autonomic remodeling and VT/VF, providing novel insight into the mechanisms underlying sympathovagal imbalance after MI.

## INTRODUCTION

Sympathoexcitation and vagal dysfunction occur after chronic myocardial infarction (MI), predispose to ventricular arrhythmias (VT/VF) and result in progression of heart failure^1,2^. Heart failure and MI have been reported to be associated with significant structural and functional pathological autonomic remodeling, including glial activation and inflammation in the peripheral autonomic ganglia^3,4^, which are thought to play a role in the sympathovagal imbalance post-MI and contribute to electrophysiological instability and VT/VF^5–8^. In addition, increased cervicothoracic spinal nociceptive afferent neurotransmission post-MI has been reported to reflexively amplify sympathetic efferent outflow^9–13^. Therefore, targeting these spinal afferent fibers/neurons, for instance, with resiniferatoxin (RTX), an ultrapotent and specific agonist of the transient receptor potential vanilloid 1 receptor (TRPV1)^14–21^, could represent an important therapeutic target. RTX ablates TRPV1-expressing nociceptive neurons via rapid and prolonged influx of calcium ions, leading to neuronal cytotoxicity, and cell death^22–28^. Consistent with this, a prior study showed that the ablation of cardiac TRPV1 neurons via direct application of RTX on thoracic dorsal root ganglia (DRG) reduced acute ischemia driven ventricular arrhythmias in a healthy porcine model^29^. Yet the effect of spinal afferent interruptions with RTX on ventricular arrhythmias has not been evaluated beyond the acute MI window, even though ventricular arrhythmias due to chronic autonomic and myocardial remodeling remain an important cause of SCD long after the acute event^1,30,31^. Hence, in this study, we first sought to assess if *chronic* spinal nociceptive ablation is an important contributor to the ensuing autonomic remodeling that leads to electrophysiological heterogeneity and ventricular arrhythmias with chronic MI. To accomplish this, we administered epidural RTX prior to MI at the C7-T1 levels to target cervicothoracic spinal TRPV1 nociceptive afferent fibers and neurons. Four-to-six weeks after chronic MI, we evaluated (i) tissue structural and neurochemical changes, (ii) functional autonomic responses, and (iii) electrophysiological stability in chronically infarcted pigs which received RTX *versus* vehicle. We next sought to evaluate the potential therapeutic value of spinal nociceptive ablation for patients with refractory VT/VF after the acute phase of MI, when chronic pathological autonomic remodeling has already developed. To that end, we tested whether acute C7-T1 epidural RTX ablation of spinal nociceptive afferents in the established post-MI setting (4-6 weeks post-MI) would be sufficient to improve autonomic balance and reduce VT/VF burden.

## RESULTS

To determine if spinal nociceptive (TRPV1) afferent signaling is an underlying contributor to pathological structural and functional autonomic remodeling after MI, these afferents were ablated by cervicothoracic (C7-T1) epidural administration of RTX (Figure 1). RTX was administered at the C7-T1 epidural level, as the majority of *cardiac* sensory innervation has been shown to pass through the C7 and upper thoracic dorsal root ganglia (DRG) and spinal horns^32^. To characterize the safety, efficacy and pharmacodynamic profile of RTX, we first administered epidural RTX to three healthy control animals. In these experiments we primarily focused on changes in heart rate (HR) and left ventricular systolic pressure (LVSP) – parameters (or their surrogates) that can be readily assessed intra-operatively in humans without the need for intraventricular catheters. After RTX administration, an initial sympathoexcitatory response was observed in theseanimals, characterized by increased HR and LVSP, followed by stabilization of both hemodynamic parameters over the next 30-60 minutes (Supplemental Figure 1). Accordingly, for the subsequent chronic studies, myocardial infarction was induced 2-3 hours after C7-T1 epidural RTX (cRTX; *n*=11) or epidural saline (vehicle; *n*=13) administration.

**Figure 1.**
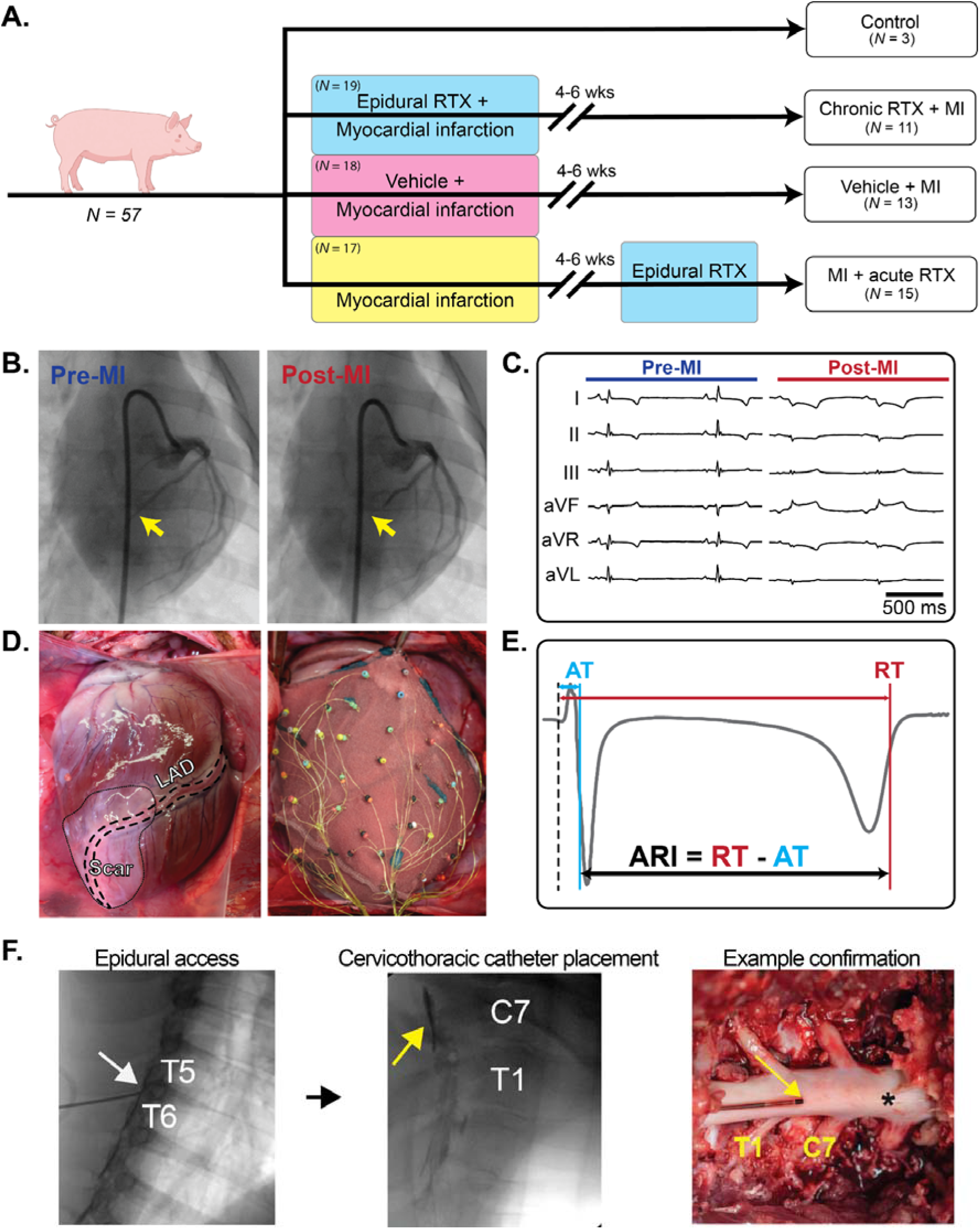
Experimental protocol. **(A)** Schematic representation of the experimental groups and their respective interventions. **(B)** Percutaneous creation of myocardial infarction (MI) via injection of microspheres in the distal left anterior descending (LAD) coronary artery after percutaneous angioplasty balloon inflation. Following injection, flow is significantly decreased in the mid and distal LAD (red arrows) and **(C)** ST-segment elevation is observed on the surface ECG. **(D)** At the time of terminal studies, a 56-electrode sock was placed around the ventricles to record local unipolar electrograms, from which **(E)** repolarization time (RT) and activation time (AT) were measured and activation recovery interval (ARI) calculated in all infarcted animals. **(F)** For epidural injections, an epidural catheter (white arrow) was advanced to the C7-T1 dorsal epidural space and position confirmed by contrast injection under fluoroscopic guidance. In terminal procedures involving the group of animals that underwent acute epidural RTX injections after chronic MI, catheter position was secondarily confirmed at the end of experiments by cut down to the epidural space and removal of epidural fat. Asterisk placed at cranial aspect of the spinal cord.

Hemodynamic profiles of animals treated with epidural RTX prior to MI (cRTX) were comparable to those of the three pilot, healthy control animals (Supplemental Figure 2), and showed that animals reached a steady hemodynamic state approximately one hour after RTX administration. Comparison of MI-related mortality in cRTX versus vehicle-treated animals showed similar survival rates between these groups (Supplemental Figure 3). However, VT/VF incidence in the acute-MI phase was higher in cRTX animals compared to vehicle-treated animals (Supplemental Figure 3), possibly due to residual sympathoexcitatory effects associated with cervicothoracic epidural RTX administration.

Finally, we sought to evaluate the pathophysiological role of nociceptive afferent signaling on subsequent functional chronic autonomic remodeling and VT. To this end, the efficacy of epidural RTX for ablating nociceptive afferents and the associated hemodynamic and autonomic responses, neural activity, and electrophysiological parameters were then assessed in both cRTX and vehicle-treated infarcted animals 4-6 weeks after MI and epidural RTX/vehicle administration were compared. We also examined the effect of RTX-mediated nociceptive neuron ablation on (i) plasma proteomic profiles and inflammatory markers, (ii) markers of glial activation and (iii) neuroinflammation in the spinal cord at both the C7-T1 as well as the lower thoracolumbar regions (T12-L1, distal anatomical control).

### Selectivity and potency of RTX for cervicothoracic spinal afferent nociceptive neurons

As most TRPV1-expressing nociceptive neurons co-express and release calcitonin gene-related peptide (CGRP)^33,34^, we used CGRP immunoreactivity (IR) as a marker for these nociceptive afferents. Since spinal nociceptive signaling is mediated by cell bodies in the DRG, which synapse on second order neurons in the upper lamina of the dorsal horn^35^, we evaluated whether administration of RTX prior to MI had resulted in chronic depletion of nociceptive fibers/neurons at these sites in cRTX (*n*=7) versus vehicle treated (*n*=5) animals (Figure 2A-D). In cRTX animals, the percentage of CGRP-IR neurons in the C7-T1 DRG was significantly decreased (14±2% *vs* 22±3%, respectively, *p*<0.05; Figure 2C), and CGRP-IR in the dorsal horn was also significantly reduced compared to vehicle animals (1.2±0.3% *vs* 5.3±0.6%, respectively, *p*<0.05; Figure 2D), confirming successful chemoablation of nociceptive afferent fibers and neurons in cRTX animals.

**Figure 2.**
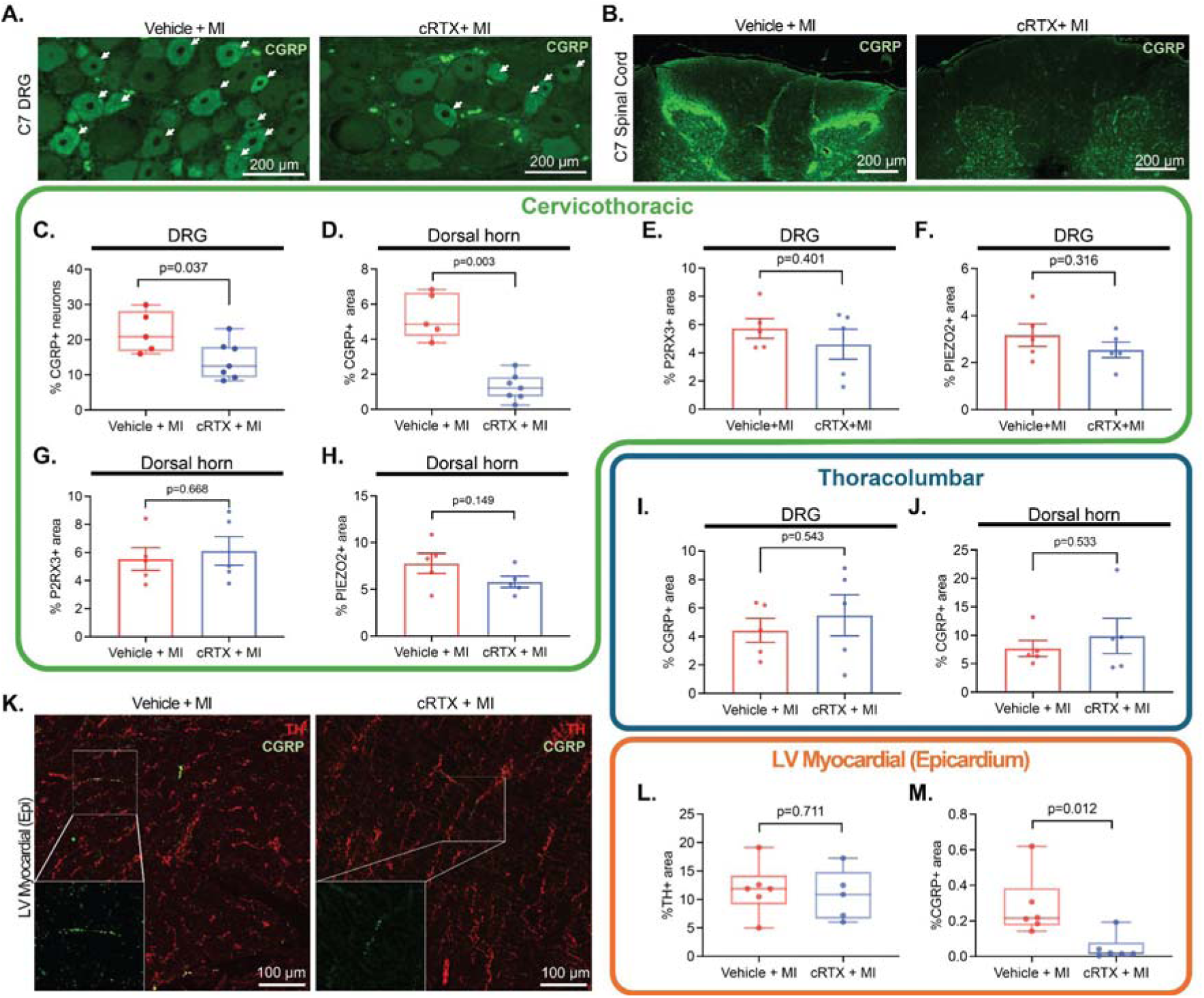
Immunohistochemical confirmation of successful ablation of cervicothoracic spinal nociceptive neurons and fibers. **(A)** Representative images of calcitonin gene-related peptide (CGRP) immunostaining of the dorsal root ganglia, and **(B)** spinal cord dorsal horn from a chronic vehicle and an epidural cRTX (chronic resiniferatoxin-treated) animal. **(C)** A significant decrease in CGRP expressing neurons in the dorsal root ganglia (DRG) and **(D)** dorsal horn of epidural cRTX (*n*=7) *vs* vehicle animals (*n*=5) is observed, confirming successful chemoablation of spinal nociceptive cardiac afferents in animals that received RTX. **(E)** Expression of purinergic receptor P2X (P2RX3) and **(F)** Piezo-type mechanosensitive ion channel component 2 (PIEZO2) in the cervicothoracic DRG was unchanged between cRTX (*n*=5) *vs* vehicle treated, infarcted animals (*n*=5). Similarly, expression levels of **(G)** P2RX3 and **(H)** PIEZO2 in the cervicothoracic dorsal horn were not different between vehicle-treated (*n*=5) versus cRTX animals (*n*=5). **(I)** Comparison of CGRP expressing neurons in the thoracolumbar (T12-L1) DRGs, or **(J)** dorsal horn of vehicle-treated (*n*=5) versus cRTX animals (*n*=5), demonstrated no difference, suggesting that these lower spinal levels were unaffected by the local (C7-T1) administration of RTX. **(K)** Representative images of tyrosine hydroxylase (TH) and CGRP immunostaining at the level of the heart. **(L)** Expression of TH (by dopaminergic non-TRPV1 expressing neurons, and sympathetic efferents) is not different in the left ventricle (LV) of RTX (*n*=5) *vs* vehicle-treated infarcted animals (*n*=5), whereas **(M)** the immunoreactivity of CGRP is significantly lower in the RTX (*n*=5) *vs* vehicle animals (*n*=5). Epicardial CGRP expression was compared using the two-sided Mann-Whitney test, all other differences between vehicle and cRTX animals were assessed using the two-sided, unpaired Student’s t-test. Epidural vehicle-treated animals that subsequently were infarcted are denoted as Vehicle+MI and epidural RTX-treated animals that were subsequently infarcted are denoted as cRTX+MI. For all bar graphs, data are presented as mean values and error bars represent SEM. For each boxplot, the center line indicates the median, the upper and lower bounds represent the first and third quartiles, and the whiskers represent the minimum and maximum values observed. Source data are provided with this paper.

Although prior literature has shown selectivity of RTX for TRPV1-expressing chemosensitive neurons^19–21^, we further confirmed the selectivity of RTX in our studies by evaluating two other major afferent populations: the chemosensitive purinergic P2RX3-expressing neurons and mechanosensitive PIEZO2-expressing neurons. Consistent with prior literature, evaluation of the C7-T1 spinal cord DRG and dorsal horns showed that the percentage of PIEZO2- and P2RX3-immunoreactive afferents was similar between cRTX and vehicle-treated animals (Figure 2E-H).

It has been shown that given the fatty nature of the epidural space, injection of anesthetics and compounds at various spinal levels, including RTX, rarely diffuse beyond 1-2 spinal levels^36–38^. However, to further confirm the anatomical and spatial selectivity of C7-T1 epidural RTX in the chronically infarcted animals, CGRP expression in the spinal cord dorsal horn and DRG at the cervicothoracic level (C7-T1) were compared to the thoracolumbar level (T12-L1). In accordance with other reports, we observed that CGRP expression was unchanged in the T12-L1 dorsal horn and DRG of vehicle-treated versus cRTX animals (Figure 2I-J).

Finally, as spinal C7-T1 afferents chiefly innervate the heart^39^, we evaluated whether ablation of nociceptive neurons at this level with RTX would manifest as a loss of CGRP-expressing fibers at the level of the ventricular myocardium. Evaluation of LV myocardium showed that the density of CGRP-expressing ventricular fibers was reduced in epidural cRTX versus vehicle treated animals (Figure 2K-M). By contrast, the abundance of tyrosine hydroxylase (TH), a marker of sympathetic *efferent* fibers, was unchanged, further underscoring the chemical selectivity of RTX for TRPV1-nociceptive neurons.

### Lack of detectable off-target effects with local epidural RTX

In line with the histological data indicating segmental selectivity in the effects of RTX, local epidural administration of RTX did not appear to cause any significant off-target effects, as no behavioral differences were noted between cRTX and vehicle-treated animals in the 4-6 weeks following MI. Furthermore, plasma levels of hepatic enzymes aspartate transaminase (AST) and alanine aminotransferase (ALT), which are clinically used to assess liver function, were similar between RTX-and vehicle-treated animals (Supplemental Figure 4). Finally, weight gain in the four-to-six-week period following vehicle or RTX treatment was similar between these groups (Supplemental Figure 5).

### Longitudinal effects of RTX prior to MI on cardiac and autonomic remodeling

Four-to-six weeks after MI, hemodynamic parameters including HR, LVSP, inotropy (dP/dt_max_), and lusitropy (dP/dt_min_) were assessed in cRTX (*n*=11) and vehicle-treated (*n*=13) animals. Baseline hemodynamic parameters were similar between these two groups of animals (Figure 3A-D), suggesting no differences in resting hemodynamic function.

**Figure 3.**
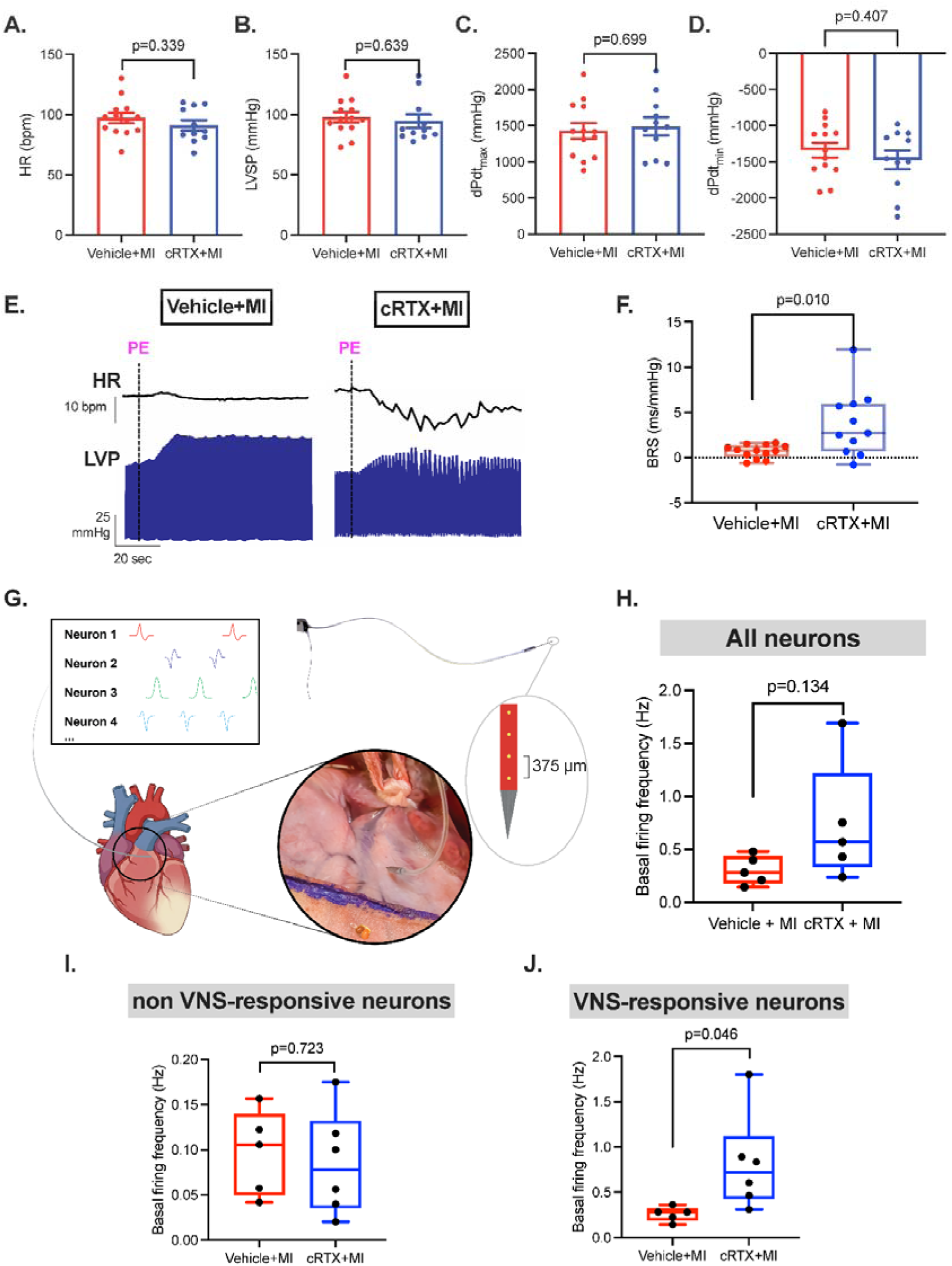
Effects of epidural RTX prior to MI on cardiac hemodynamics and vagal tone four to six weeks after MI. **(A-D)** Baseline hemodynamic parameters were not different between vehicle (*n*=13) and epidural cRTX (*n*=11) animals. **(E)** Example of phenylephrine-evoked baroreflex responses in a vehicle and epidural cRTX (resiniferatoxin-treated, infarcted) animal. **(F)** Baroreflex sensitivity (BRS) was significantly improved in epidural cRTX (*n*=11) compared to vehicle animals (*n*=13). **(G)** Schematic image of ventral interventricular ganglionated plexus (VIVGP) neural recordings and identification of VIVGP neurons. A magnified image of the neural recording electrode in a VIVGP is shown. The left atrial appendage is being retracted to fully expose the VIVGP. A picture of the neural probe and a schematic of the tip with the most distal 4 recording electrodes are shown. There was no difference in basal firing frequency of all **(H)** or non-VNS-responsive **(I)** neurons. **(J)** However, the basal firing frequency of VNS-responsive neurons (*e.g.*, parasympathetic postganglionic neurons) was significantly higher in epidural cRTX (*n*=6) compared to vehicle animals (*n*=5). Differences in heart rate (HR), left ventricular systolic pressure (LVSP), inotropy (dP/dt_max_), lusitropy (dP/dt_min_), and neuronal firing between vehicle and epidural cRTX animals were assessed using the two-sided unpaired Student’s *t*-test, and BRS was compared by the two-sided Mann-Whitney test. For all bar graphs, data are presented as mean values and error bars represent SEM. For each boxplot, the center line indicates the median, the upper and lower bounds represent the first and third quartiles, and the whiskers represent the minimum and maximum values observed. PE: Phenylephrine. Epidural vehicle-treated animals that subsequently were infarcted are denoted as Vehicle+MI and epidural RTX-treated animals that were subsequently infarcted are denoted as cRTX+MI. Source data are provided with this paper.

We then assessed vagal baroreflex sensitivity (BRS), a marker of parasympathetic function^40,41^, by evaluating responses to infusion of phenylephrine (an alpha-1 adrenergic agonist). BRS was significantly increased in cRTX (*n*=11) compared to vehicle animals (*n*=13; 3.7±1.1 *vs* 0.7±0.2 ms/mmHg, respectively, *p*<0.05; Figure 3E-F). These findings support the key contribution of spinal nociceptive afferents to post-MI vagal baroreflex impairment, which has been linked to increased SCD risk^40,42,43^.

To further confirm the impact of sympathetic spinal nociceptive afferent signaling on *efferent* vagal dysfunction post-MI, activity of intra-cardiac, postganglionic parasympathetic neurons in the ventral interventricular ganglionated plexus (VIVGP), a major source of ventricular parasympathetic innervation^44–46^, were measured in cRTX (*n*=6) versus vehicle (*n*=5; Figure 3G) animals. We defined post-ganglionic parasympathetic neurons as neurons exhibiting statistically significant alterations in their vagal activity in response to low level cervical vagus nerve stimulation (VNS)^47,48^. Low level cervical VNS permits the assessment of neuronal activity without confounding hemodynamic changes^47–49^, enabling the classification and evaluation of neurons based on whether or not they receive vagal input. Spinal nociceptive afferent blockade prior to MI (cRTX animals) neither affected overall bulk firing of VIVGP neurons (Figure 3H) nor the firing of VNS non-responsive neurons (Figure 3I). By contrast, the baseline firing of VNS-responsive neurons (i.e., post-ganglionic cardiac parasympathetic neurons) was significantly higher in cRTX versus vehicle animals (Figure 3J). Together with the BRS findings, these data suggest that interruption of cervicothoracic spinal afferents prior to MI preserved vagal function after chronic MI, implicating nociceptive afferent neurons as contributors to the reduction in vagal tone observed after MI.

Lastly, we tested reflex-driven autonomic responses by the application of bradykinin and capsaicin to the ventricular epicardium to activate cardiac nociceptive afferent nerve endings. Epicardial bradykinin elicited primarily a sympathoexcitatory response in vehicle-treated animals (*n*=10), whereas cRTX animals (*n*=11) exhibited a predominantly vagal response (decreased HR and LVSP; *p<*0.05 for all; Figure 4A-B). Because sympathetic stimulation can increase dispersion of repolarization (DoR)^50^, we further assessed ventricular electrophysiology and found that, in cRTX animals, bradykinin application prolonged repolarization time and reduced DoR (*p<*0.05; Figure 4C-D), consistent with improved electrophysiological stability.

**Figure 4.**
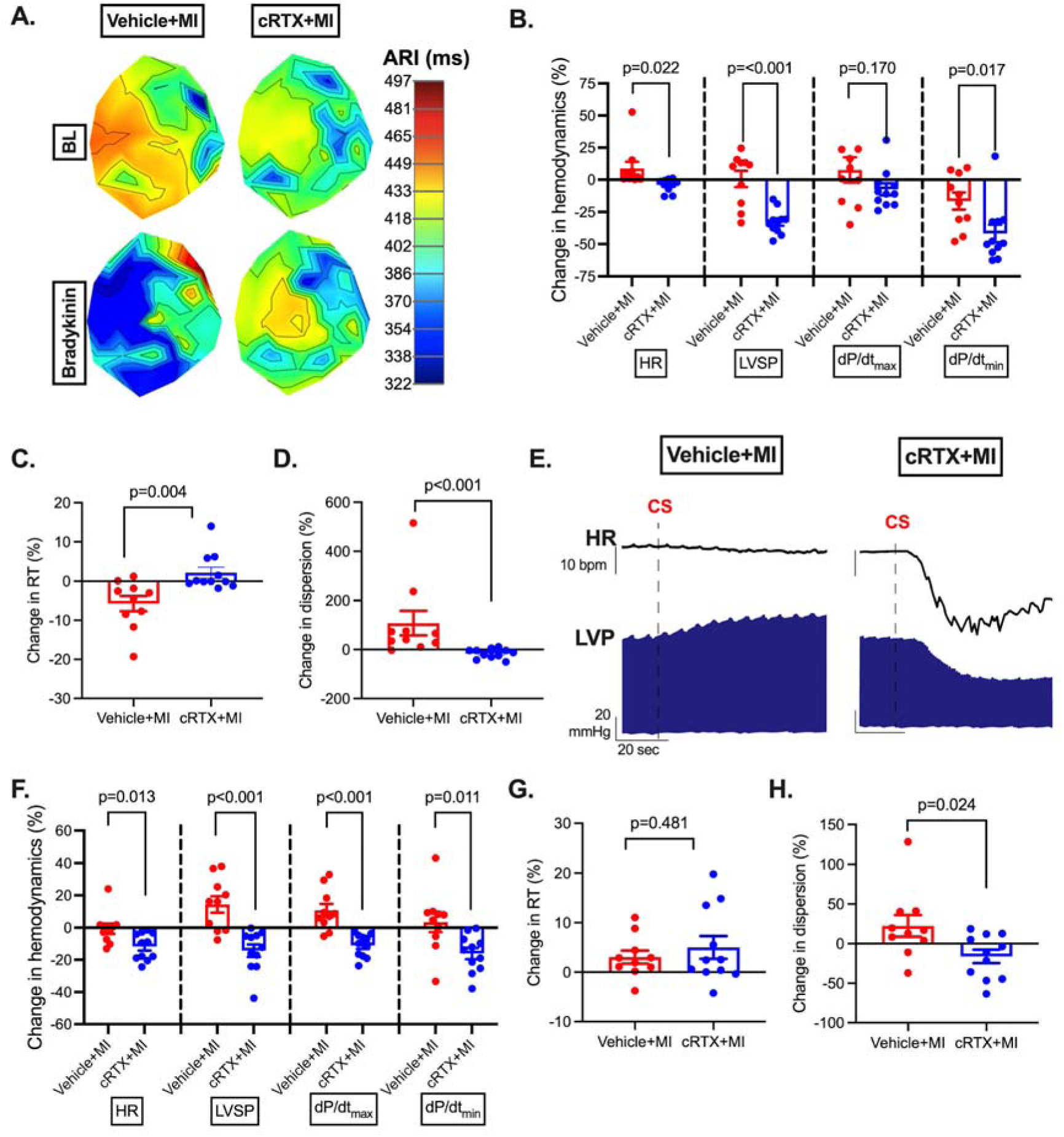
Effects of spinal afferent ablation with epidural RTX prior to MI on reflex-evoked cardiac function four-to-six weeks after MI. **(A)** Representative 2D-map of electrophysiological changes evoked by epicardial application of bradykinin in a vehicle (left) and an epidural cRTX (RTX-treated, infarcted) animal (right). **(B-D)** Quantified hemodynamic and electrophysiological responses to epicardial application of bradykinin in vehicle-treated (*n*=10) and cRTX (*n*=11) infarcted animals are shown. **(E)** Representation of hemodynamic changes evoked by epicardial application of capsaicin. **(F-H)** Quantified hemodynamic and electrophysiological changes in repolarization time (RT) and dispersion of RT upon epicardial application of capsaicin in vehicle (*n*=10) and cRTX (*n*=11) animals. Differences in heart rate (HR), left ventricular systolic pressure (LVSP), dP/dt_max_, dP/dt_min_, RT and dispersion responses upon bradykinin and capsaicin administration between vehicle and epidural cRTX animals were assessed using the two-sided unpaired Student’s *t*-test. CS: Capsaicin. Epidural vehicle-treated animals that subsequently were infarcted are denoted as Vehicle+MI and epidural RTX-treated animals that were subsequently infarcted are denoted as cRTX+MI. For all bar graphs, data are presented as mean values and error bars represent SEM. MI = myocardial infarction.

Like bradykinin, capsaicin increased LVSP, LV inotropy, and DoR in vehicle-treated animals, consistent with a predominantly sympathetic and pro-arrhythmic response. However, in animals that had received epidural RTX prior to MI (cRTX animals), these effects were reversed at 4-6 weeks post-MI, with capsaicin evoking a vagal-predominant response with reduced observed DoR, suggestive of reduced arrhythmia susceptibility (Figure 4E-H).

### RTX therapy prior to MI impedes post-MI oxidative stress, inflammation, and glial activation

Cardiovascular disease is characterized by systemic inflammatory responses and oxidative stress^51–53^. After MI, heightened sympathetic outflow amplifies inflammatory tone by noradrenergic signaling to hematopoietic niches, mobilizing myeloid cells into the circulation^54^. Thus, to assess if spinal nociceptive afferent signaling may also contribute to this post-MI inflammatory phenotype, we assayed differences in the plasma proteome by ELISA (Supplemental Figure 6) and mass spectrometry of cRTX (*n*=10) and vehicle-treated (*n*=9) animals (Figure 5A). Differentially expressed proteins in epidural cRTX versus vehicle animals mapped chiefly to immune response, oxidative stress, and cell cycle pathways (Figure 5B; Supplementary data 1). Notably, PSMD8, ZAP70 and VAV1, key immune regulators^55–58^, were downregulated four-to-six weeks after MI in cRTX animals. Moreover, pathways associated with immune activation and oxidative stress were also diminished in cRTX animals (Figure 5C; Supplementary data 2). Conversely, the ERK1/2 cascade which has been reported to be fundamental for cardiomyocyte homeostasis^59^, was significantly increased in cRTX animals. Mean plasma levels of TNF-alpha at 4-6 weeks post-MI were significantly reduced in cRTX animals, while mean IL-12 plasma levels, although different, did not reach statistical significance (*p*=0.052, Supplemental Figure 6). Levels of IL-6 and IL-10 were below detection for both groups.

**Figure 5.**
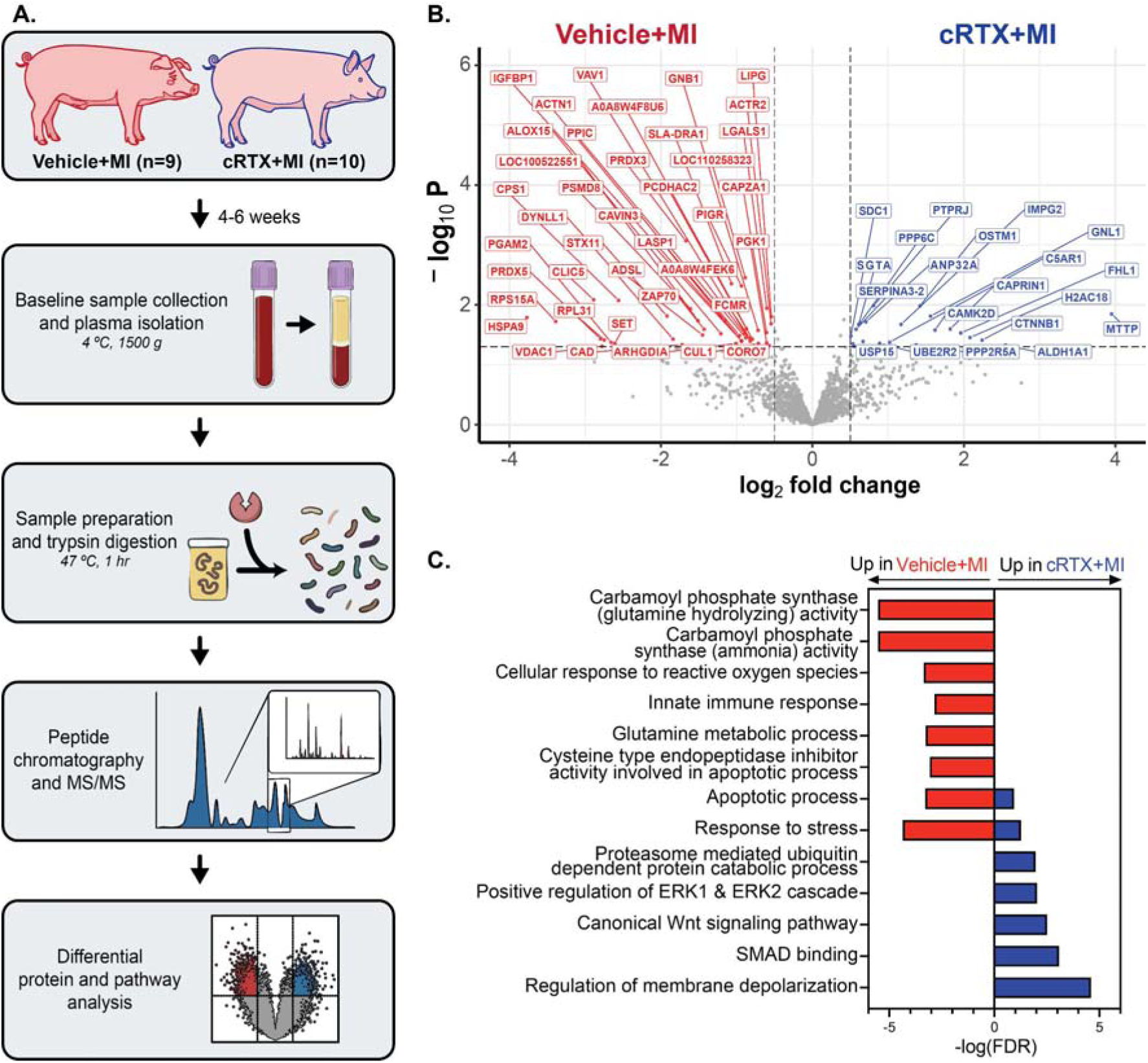
Proteomic analyses demonstrate a decrease in inflammatory and stress response pathways in infarcted animals with epidural RTX treatment prior to MI compared to untreated, infarcted animals four-to-six weeks post infarction. **(A)** Schematic representation of plasma protein isolation to proteomic data analyses. **(B)** Volcano plot showing the differentially expressed proteins in vehicle and cRTX (resiniferatoxin-treated, infarcted) animals. **(C)** Proteomic pathway analyses conducted on expression data against the Gene Ontology database show a significant decrease in oxidative stress and inflammatory pathways in cRTX infarcted animals compared to vehicle-treated, infarcted animals. Pathways associated with extracellular signal-regulated kinase 1 (ERK1) and extracellular signal-regulated kinase 2 (ERK2) cascades were upregulated in cRTX animals, which have been described to be fundamental for cardiomyocyte homeostasis and stress responses. Differentially expressed proteins were identified using a two-sided Student’s t-test. Pathway analysis was performed against the Gene Ontology database using Rapid Integration of Term Annotation and Network. Animals that received vehicle (epidural saline) prior to MI are denoted as Vehicle+MI and animals that received epidural RTX prior to MI are denoted as cRTX+MI; *n*=9 for Vehicle+MI, *n*=10 for cRTX+MI. MS = mass spectrometry. Source data are provided with this paper.

Given these effects on the systemic inflammatory response, we next asked whether spinal afferent signaling contributes to local spinal neuroinflammation and glial activation in the chronic post-MI period. To this end, we also quantified immunoreactivity for activated satellite glial cells (GFAP), microglia (IBA1), and T-cells (CD3) in the C7 and T1 dorsal horns of cRTX (*n*=7) versus vehicle (*n*=5). cRTX, infarcted animals showed reductions in GFAP, IBA1, and CD3 versus vehicle-treated controls (Figure 6A-G).

**Figure 6.**
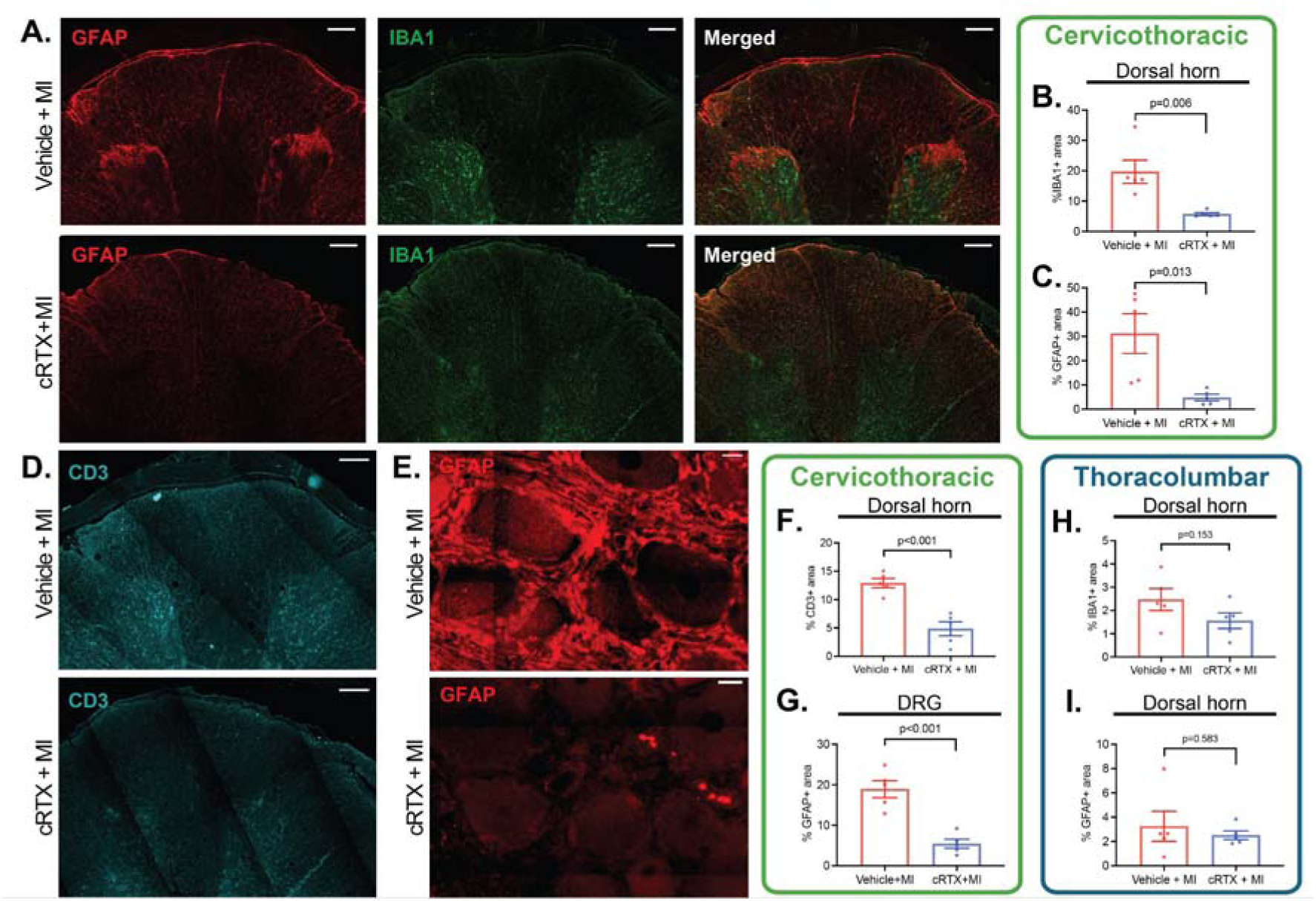
Immunohistochemical changes associated with ablation of cervicothoracic spinal nociceptive nerves with epidural RTX. **(A)** Representative images of immunostaining of spinal horns in cRTX and vehicle animals for glial fibrillary acidic protein (GFAP) and ionized calcium-binding adaptor molecule 1 (IBA1; markers of glial activation and microglia, respectively (scale bars represent 500 µm). **(B-C)** Quantified data show decreased IBA1 **(B)** and GFAP **(C)** immunoreactivity in RTX-treated (cRTX; *n*=5) *vs* vehicle-treated (*n*=5) infarcted animals. **(D)** Representative images of immunostaining of the spinal horn for CD3 (marker of T lymphocytes). **(E)** Representative image of immunohistochemical staining of GFAP in T1 (thoraic level 1) dorsal root ganglia (scale bars represent 20 µm). **(F)** Quantified data show a significant reduction in the amount of CD3 positive T lymphocytes in cRTX (*n*=5) *vs* vehicle-treated (*n*=5) infarcted animals, suggesting reduced inflammation. **(G)** Quantified data showed a significant decrease in glial activation in the high thoracic dorsal root ganglia of cRTX (*n*=5) versus vehicle (*n*=5) animals. These effects of epidural RTX administration on reducing inflammation and glial activation were localized to the high thoracic region, as spinal cord dorsal horn expression of **(H)** IBA1, and **(I)** GFAP in the thoracolumbar region were similar between the vehicle-treated (*n*=5) versus cRTX (*n*=5). Differences between vehicle-and cRTX animals were assessed using the two-sided unpaired t-test. Animals that received vehicle (epidural saline) prior to MI are denoted as Vehicle+MI, and animals that received epidural RTX prior to MI are denoted as cRTX+MI. For all bar graphs, data are presented as mean values and error bars represent SEM. Source data are provided with this paper.

Finally, we evaluated whether GFAP and IBA1 immunoreactivity were different at the level of the T12-L1 spinal horns. There was no difference in GFAP-IR or IBA1-IR in cRTX (n=5) versus vehicle-treated (n=5) infarcted animals in the thoracolumbar dorsal horns (Figure 6H-I). Notably, vehicle treated animals showing decreased GFAP and IBA1 in the thoracolumbar region as compared to their cervicothoracic region, suggesting that increased neuroinflammation in the cervicothoracic region is driven by the visceral organ affected (i.e. the heart), consistent with prior literature showing increased neuronal cFOS in the spinal cord limited to upper thoracic regions after MI^60^.

### Effects of RTX prior to MI on scar size, ventricular electrophysiology and arrhythmias

Scar size and detailed electrophysiological measurements were evaluated in RTX (*n*=11) versus vehicle treated infarcted animals (*n*=13) to determine if epidural RTX prior to MI had altered these after chronic MI. Four-to-six weeks after MI, the size of ventricular scars were not different between those that had received RTX versus vehicle prior to MI (Supplemental Figure 7), as assessed by epicardial LV and RV voltage mapping and quantification of low voltage areas, defined as border zone and scar regions (percent low voltage area: 23.7±2.3% in cRTX (*n*=11) *vs* 22.3±1.9% in vehicle (*n*=13) animals, *p*=0.634) and gross pathological examination and image quantification of dense scar regions of explanted hearts (scar size: 10.0±1.3 % *vs* 10.7±1.3% of anterior ventricular surface in cRTX (*n*=11) *vs* vehicle (*n*=13) animals, respectively, *p*=0.690). In addition, analysis of echocardiographic data showed that LV end systolic and end-diastolic areas were not significantly different between the two groups, with a similar change in these dimensions (quantified as LV ejection fraction or % change in systole) of 40.0±2.8% *vs* 36.0±3.8% in cRTX (*n*=5) *vs* vehicle-treated (*n*=5) animals, respectively, *p*=0.540).

For regional electrophysiological analyses, ARIs were compared between scar, border zone, and viable regions, with electrodes selected from each area based on epicardial voltage mapping^49,61^. Baseline (resting) global ventricular ARIs and regional ARIs were not significantly different between infarcted epidural cRTX-versus vehicle-treated animals (Figure 7A-C). Moreover, basal ventricular global DoR was not significantly different between these groups (Figure 7D). However, granular regional analyses demonstrated that the border zone dispersion was significantly greater in vehicle vs. cRTX animals (984±234 *vs* 351±154 ms^2^, respectively, *p*<0.05; Figure 7E). While atrial effective refractory period (ERP) was not significantly different (Figure 7F), ventricular endocardial ERP (measured at the right ventricular apex) was significantly longer in cRTX versus vehicle animals (283±5 *vs* 267±5 msec, respectively, *p*<0.05, Figure 7G-H), suggesting increased ventricular refractoriness due to ablation of spinal nociceptive neurons/fibers. Finally, DoR of the prematurely paced beat (S2), which is known to activate spinal receptors and reflexively increase sympathetic outflow resulting in ventricular arrhythmias^62,63^, was significantly less in epidural cRTX (cRTX: 817±68 msec^2^) *vs* vehicle animals (1163±90 msec^2^, *p*<0.05, Figure 7I). Importantly, cRTX animals were less inducible for VT/VF with extra-stimulus pacing (18% in cRTX *vs* 62% in vehicle, *p*<0.05; Figure 7J-L), while there was no difference in pacing thresholds to capture the ventricular myocardium (Supplemental Figure 8).

**Figure 7.**
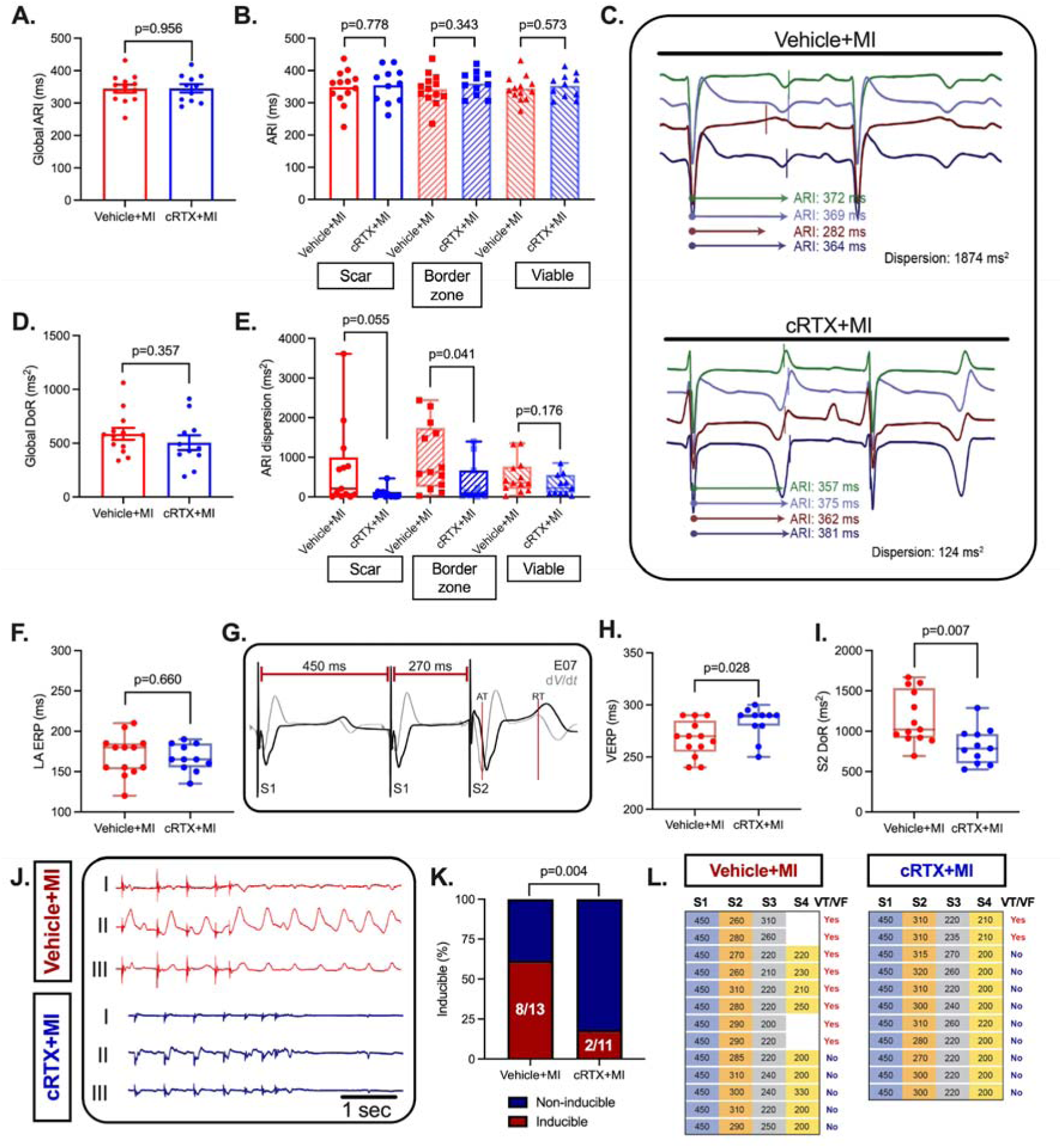
Effects of spinal nociceptive afferent ablation prior to MI on ventricular APD, DoR, refractoriness, and ventricular arrhythmia inducibility post-infarction. **(A)** Administration of cervicothoracic epidural RTX prior to MI did not alter global baseline activation recovery intervals (ARI), or **(B)** regional ARIs four-to-six weeks post-MI. **(C)** Examples of border zone electrograms from a vehicle (top) and RTX treated (cRTX+MI, bottom) animal showing ARIs from these regions. **(D)** In cRTX animals, global dispersion of repolarization (DoR) was not significantly different from vehicle-treated infarcted animals, however, **(E)** regional analyses demonstrated that ARI dispersion was greatest in border zone regions in both vehicle and RTX treated animals, but that this border zone dispersion was significantly less in infarcted animals treated with RTX. **(F)** Left atrial ERP (LA ERP) was not significantly different between vehicle-treated and cRTX animals. **(G)** Example of ventricular ERP (VERP) measurements showing the last-captured extra-stimulus (S2) at interval of 270 msec. **(H)** Ventricular ERP (VERP) was significantly prolonged in RTX treated animals, and **(I)** RT-dispersion of the S2 paced beat was significantly decreased in cRTX animals. **(J)** Example of VT/VF induction using programmed stimulation. The epidural vehicle-treated MI animal (upper panel) was inducible with 1 extra-stimulus, but the cRTX animal (lower panel) was not inducible. **(K)** VT/VF inducibility was reduced from 62% in vehicle-treated infarcted animals to 18% in cRTX animals with up to four extra-stimuli. **(L)** Breakdown of stimulation parameters used for induction of VT/VF. All electrophysiological parameters were compared between vehicle and cRTX animals using the two-sided unpaired Student’s t-test, VT inducibility was compared by the two-sided exact binomial test. *n*=13 for vehicle and *n*=11 for epidural cRTX for all reported electrophysiological parameters. Animals that received vehicle (epidural saline) prior to MI are denoted as Vehicle+MI, and animals that received epidural RTX prior to MI are denoted as cRTX+MI. For all bar graphs, data are presented as mean values and error bars represent SEM. For each boxplot, the center line indicates the median, the upper and lower bounds represent the first and third quartiles, and the whiskers represent the minimum and maximum values observed. Source data are provided with this paper.

### RTX administration after established infarction acutely rescues arrhythmogenic phenotype

Having shown that administration of epidural RTX and ablation of cervicothoracic nociceptive neurons prior to MI (cRTX) impeded autonomic/electrophysiological remodeling and reduced VT/VF inducibility at 4-6 weeks post-MI, we next asked whether RTX could still be therapeutic benefit after pathological remodeling has been established. To test this hypothesis, a separate group of pigs underwent myocardial infarcts, and 4-6 weeks later, received cervicothoracic epidural RTX (aRTX, *n*=15); hemodynamic and autonomic responses were assessed acutely every hour up to 4 hours after RTX administration.

Consistent with animals receiving RTX prior to MI (cRTX), aRTX induced an initial sympathoexcitatory response with peak increases in HR (84±4 to 87±4 beats/min, *p=*0.001), LVSP (118±5 to 131±7 mmHg, *p*<0.01), and inotropy (1393±67 to 1484±79 mmHg/s, *p*<0.01) within the first hour, Figure 8. By 4 hours after epidural RTX administration, HR had stabilized but remained higher than pre-RTX values, while LV inotropy and LVSP were slightly lower (Figure 8A-E).

**Figure 8.**
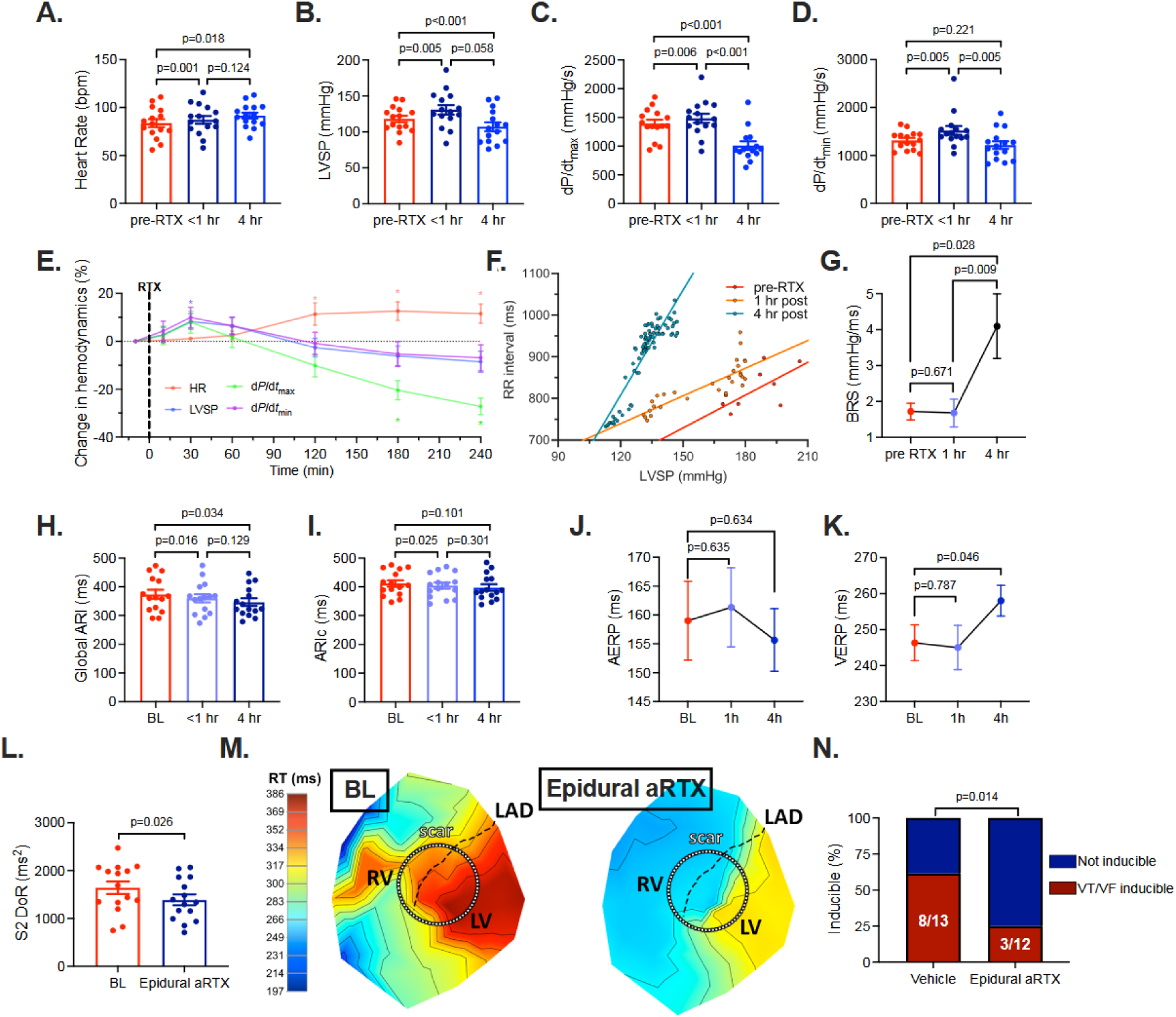
Temporal hemodynamic responses and electrophysiological effects of acute cervicothoracic spinal nociceptive afferent ablation with epidural RTX in the setting of chronic MI. The acute effects of epidural RTX (aRTX) were assessed to determine the kinetics and tolerability of RTX in infarcted animals (*n*=15). Within 1-hour of administration, RTX increased **(A)** heart rate (HR), **(B)** left ventricular systolic pressure (LVSP), **(C)** left ventricular inotropy (dP/dt_max_), and **(D)** left ventricular lusitropy (dP/dt_min_). Left ventricular (LV) lusitropy remained significantly elevated, whereas LV inotropy and LVSP were decreased 4 hours after epidural RTX administration. **(E)** Percentage change in HR, LVSP, LV inotropy and lusitropy over time are shown. *p < 0.05 vs baseline. **(F)** Beat-to-beat changes in RR interval *vs* LVSP were plotted to assess baroreflex sensitivity (BRS) before, 1-hour, and 4-hours post-RTX injection. **(G)** BRS was significantly improved by 4-hours post-RTX (*n*=10). **(H)** Acute epidural RTX shortened activation recovery intervals (ARI) at 1- and 4-hours after administration. **(I)** When corrected for heart rate (ARIc), this effect was no longer significant. **(J)** Epidural RTX did not alter atrial ERP (AERP) in chronic MI animals. **(K)** Epidural RTX prolonged VERP 4-hours after administration. **(L-M)** This was accompanied by a reduction in the S2 dispersion of repolarization time (DoR). **(N)** While 8/13 vehicle animals were inducible for sustained VT/VF, only 3/12 epidural aRTX animals were inducible at 4 hours post-RTX administration. Serial hemodynamic and electrophysiological parameters were compared by repeated-measures ANOVA. Pre-RTX vs. <1 hour or 4 hour and 1 hour vs. 4 hour comparisons were performed only if repeated measure ANOVA was statistically significant and compared using two sided paired Student’s t-test (uncorrected). Serial BRS was compared using the Friedman test (uncorrected) and RT dispersion was compared using a two-sided paired Student’s t-test. Inducibility was compared using the two-sided exact binomial test. BL= Baseline, RV = Right ventricle, LV = Left ventricle, LAD = Left anterior descending artery. For all bar graphs, data are presented as mean values and error bars represent SEM.

While hemodynamic parameters peaked within the first hour after RTX, no effect on vagal function, as assessed by BRS, was observed at one hour (from 1.8±0.2 pre-RTX to 1.7±0.4 ms/mmHg 1-hour post-RTX, *p*>0.05; *n*=10, Figure 8F-G). By four hours after aRTX, however, BRS had significantly increased to 3.9±0.7 ms/mmHg (*p*<0.05), suggestive of augmented vagal function (Figure 8F-G).

While previous studies have acutely demonstrated the clinical anti-arrhythmic efficacy of thoracic epidural anesthesia using lidocaine or bupivacaine^64,65^, the differential contributions of efferent versus afferent blockade to ventricular arrhythmias remains unclear. Furthermore, whether acute spinal afferent blockade alone could be anti-arrhythmic is unknown. Hence, the effects of spinal sympathetic afferent nociceptive ablation on cardiac electrical stability were evaluated before and four hours after epidural RTX administration in chronically infarcted animals. After epidural aRTX, global ventricular ARIs significantly shortened (374±16 pre-RTX to 345±14 msec post-RTX; *p*<0.05, Figure 8H), whereas heart rate-corrected ARI (ARIc) was unchanged, indicating a HR-dependent effect, Figure 8I. Atrial ERP remained unchanged after aRTX (*p*>0.05; Figure 8J), but right ventricular endocardial ERP significantly increased from 246±5 pre-RTX to 258±4 msec four hours post-RTX (*p*<0.05), Figure 8K. Moreover, while epidural aRTX did not alter DoR during sinus rhythm, it significantly decreased DoR of the ventricular extra-stimulus (S2) paced beat (1639±130 pre-RTX *vs* 1389±112 msec^2^ post-RTX, *p*<0.05, Figure 8L-M). No animals developed spontaneous ventricular arrhythmia following administration of epidural RTX, despite the initial sympathoexcitatory response and presence of a chronic infarct. However, at 4 hours post-RTX administration, only 3 of 12 infarcted animals were inducible for VT/VF, significantly fewer than the chronic infarcted animals who had only been treated with vehicle (25% *vs* 62%, respectively; *p*<0.05; Figure 8N).

## DISCUSSION

The present study demonstrates that cervicothoracic spinal nociceptive afferent signaling is a critical factor in initiating and sustaining the chronic pathological autonomic remodeling after MI (Figure 9). Given the multifactorial etiology of ventricular arrhythmias, we elected to dissect the role of spinal nociceptive afferent signaling on autonomic, inflammatory, and electrophysiological mechanisms underlying the increased risk of ventricular arrhythmias after chronic MI. Local interruption of spinal TRPV1 afferent signaling after MI impeded the development of vagal dysfunction, suppressed neuroinflammation and glial activation, attenuated systemic inflammation and oxidative stress, and stabilized cardiac electrophysiological parameters, reducing VT/VF inducibility. These results highlight the potentially critical role of spinal TRPV1-expressing afferents in the development of the multitude of chronic pathological autonomic remodeling processes that have been described after MI^1–3,52,66–69^ and provide a potential target for ameliorating these effects. Importantly, even when delivered after established MI and pathological remodeling (4-6 weeks post-MI), RTX-mediated ablation was sufficient to enhance vagal tone and mitigate electrophysiological instability, corresponding to fewer inducible ventricular arrhythmias. Collectively, these findings highlight the role of spinal afferents in contributing to the development of autonomic remodeling after MI as well as support their role in sustaining autonomic dysfunction and pro-arrhythmia.

**Figure 9.**
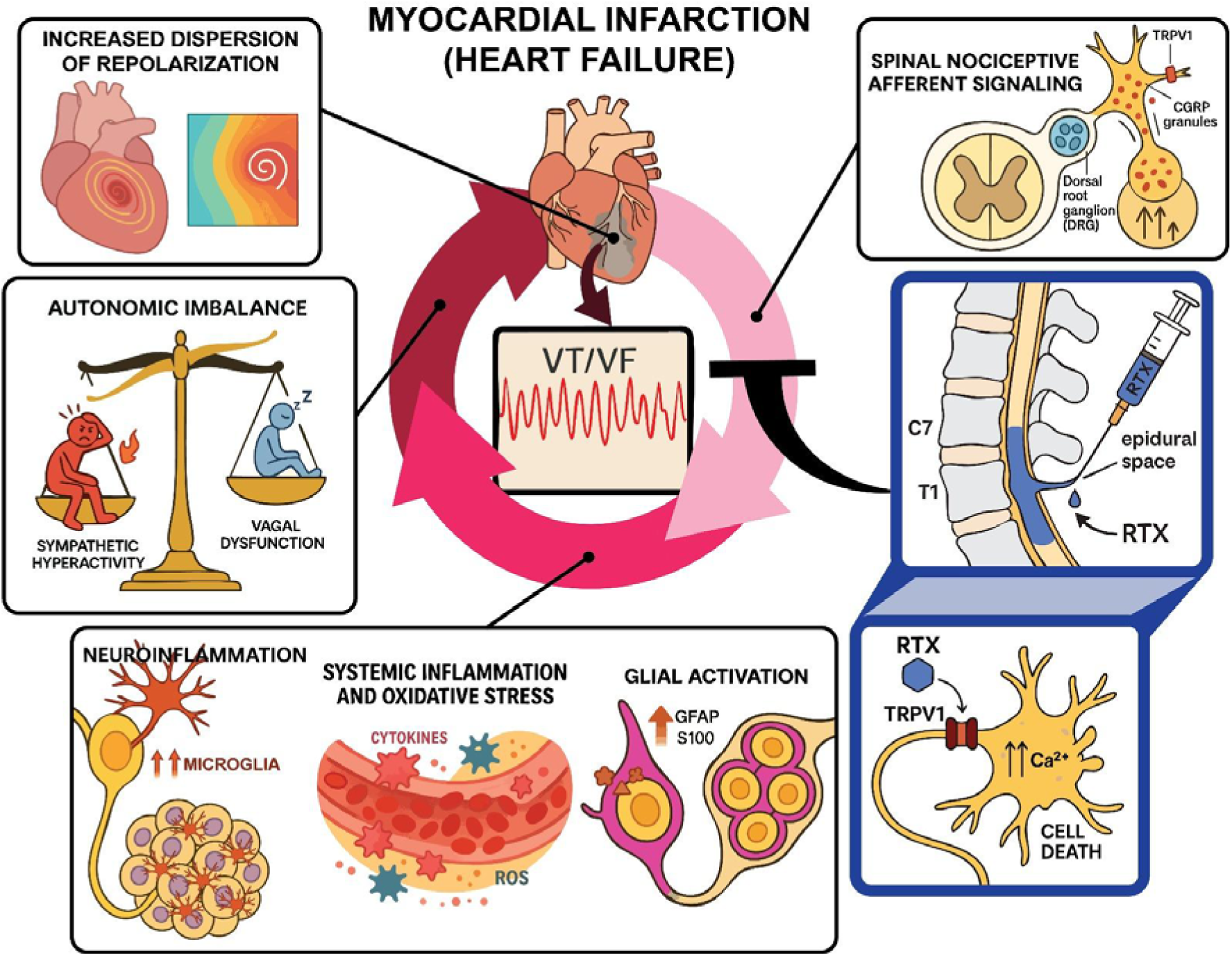
Spinal nociceptive afferent signaling contributes to post-myocardial infarction neuroinflammation, oxidative stress, glial activation, and sympatho-vagal imbalance, resulting in increased dispersion of repolarization and ventricular arrhythmias (VT/VF), a process that is mitigated by interruption of this signaling with epidural resiniferatoxin (RTX). GFAP = glial fibrillatory acid protein, TRPV1 = transient receptor potential vanilloid 1, CGRP = calcitonin gene-related peptide.

These autonomic changes merit closer consideration. After MI, pathological autonomic remodeling culminates in sympathoexcitation and vagal withdrawal, as reflected in decreased baroreflex sensitivity, heart rate variability, and altered activity of post-ganglionic parasympathetic neurons^1–3,52,66–69^, ultimately contributing to VT/VF. Reduced vagal function, as assessed by BRS, is an independent predictor of SCD and VT/VF in patients with MI and heart failure^40,42^. Persistent chronic cervicothoracic spinal afferent signaling after MI results in increased efferent sympathetic outflow. However, it has also been reported to contribute to vagal dysfunction, as suggested by experiments where activation of these fibers resulted in reduced BRS and, correspondingly, the improvement in BRS and cardiac post-ganglionic vagal neuronal activity observed with thoracic epidural anesthesia^10,48,70^.

In the present study, ablation of nociceptive TRPV1-expressing neurons prior to MI (cRTX) preserved vagal function four-to-six weeks after MI. Moreover, responses to epicardial application of either the direct TRPV1 agonist capsaicin^71^ or the B2 receptor ligand bradykinin, which sensitizes TRPV1+ neurons at the time of myocardial ischemia,^72^ were blunted in RTX-treated MI animals. In fact, while vehicle-treated infarcted animals exhibited an overt sympathoexcitatory response, cRTX animals exhibited a predominantly vagal response, accompanied by reductions in DoR, consistent with greater electrical stability. Following MI, denervation and heterogeneous nerve sprouting at scar/border zone regions is reported^5,73,74^, further amplifying electrical heterogeneity during sympathetic activation, increasing DoR and predisposing to VT/VF^75–78^. The decrease in DoR is also consistent with the potentially cardioprotective effects of vagal TRPV1-positive afferents^79,80^ that do not pass through the dorsal root ganglia or the epidural space, and therefore, remain intact.

These functional changes are paralleled by remodeling at the cellular level. MI and heart failure with reduced ejection fraction have been shown to induce satellite glial cell activation in the peripheral autonomic ganglia, spinal cord, and autonomic regions of the brain^3,66,81–86^. Satellite glial cells have been reported to respond to increased neuronal activity but may themselves augment neuronal synaptic transmissions^87^, potentially sustaining increased sympathetic reflexes. Importantly, the interplay of these cells has been shown in the stellate ganglia to establish a feed-forward process which compounds sympathetic activity^88^. In the present study, four-to-six weeks after MI, we observed diminished levels of glial activation in both the spinal cord and DRG of animals treated with RTX prior to MI. However, as satellite glial cells do not express TRPV1, this decrease may be secondary to diminished neuronal afferent activation^89,90^. The concomitant attenuation of afferent signaling and local satellite glial cell activation may cumulatively blunt efferent neuronal activation and reflex activity through the spinal cord^87^, thereby improving sympathovagal balance. Taken together, preservation of parasympathetic tone, coupled with reduced molecular and cellular markers of autonomic remodeling in the peripheral ganglia of RTX-treated animals support the hypothesis that spinal nociceptive afferents play a critical role in the pathological autonomic remodeling observed in chronic MI.

Beyond these autonomic effects, MI is associated with systemic inflammation, oxidative stress^52,53,81,91,92^, and neuroinflammation^93,94^. Elevated inflammatory cytokines^95–100^ and increased oxidative stress^101–105^ are associated with poorer outcomes and an increased risk for VT/VF in patients with MI and heart failure. However, clinical trials of anti-inflammatory therapies (e.g., TNF-〈, IL-1β, and IL-6 inhibitors) have yielded mixed results in this patient population^106–108^. In this study, plasma proteomics indicated lower markers of inflammation and stress in cRTX versus vehicle-treated animals. In parallel, we also observed reductions for microglial and T lymphocyte markers in the cervicothoracic dorsal horn of epidural RTX treated animals, suggesting that RTX treatment attenuated chronic neuroinflammation. It is thus possible that spinal nociceptive afferent signaling may contribute to the heightened neuroinflammatory milieu observed after MI. Furthermore, beyond reducing local neuroinflammation, the improved vagal tone after RTX treatment may also engage the splenic cholinergic anti-inflammatory reflex^109^, where acetylcholine binds to nicotinic receptors on white blood cells, to attenuate systemic cytokine release and systemic inflammation^110–112^.

These inflammatory and autonomic processes converge on the electrophysiological substrate for arrhythmia. Acute MI causes myocardial injury, axonal denervation, and localized nerve sprouting^68,113^. In the setting of sympathoexcitation, these changes culminate in electrical heterogeneity, especially in border zone regions, serving as the substrate for VT/VF^8,114,115^. In the present study, treatment with cervicothoracic epidural RTX prior to MI (cRTX) led to a reduction in DoR (i.e., electrical heterogeneity) in border zone myocardium four-to-six weeks post-MI, consistent with reduced electrical heterogeneity. Border zone regions, due to their innervation and myocardial heterogeneity, serve as sites for both the trigger and substrate of VT/VF^116–118^. Interestingly, the reductions in VT/VF-inducibility in RTX-treated animals were comparable to the anti-arrhythmic benefits reported with thoracic epidural anesthesia using bupivacaine^64^ and cardiac sympathetic denervation in animal models post-MI and in patients with refractory VT/VF^119,120^, interventions that simultaneously interrupt all spinal afferent *and* efferent sympathetic signaling. Given that RTX selectively targets nociceptive afferents in particular, the similar magnitude of anti-arrhythmic effects raises the possibility that a substantial portion of thoracic epidural anesthesia’s benefit is derived specifically from the blockade of these fibers.

Having established these effects when ablation preceded infarction, we next considered whether spinal afferent signaling also contributes once chronic scar and autonomic remodeling are already established. A prior study in pigs without a myocardial infarction assessed the effects of T2-T4 DRG RTX application 30 minutes prior to induction of ischemia on VT/VF burden during the acute ischemia and one hour reperfusion period^29^. In light of that report and the observed, longitudinal effects of RTX prior to MI reported herein, we subsequently aimed to evaluate the effects of C7-T1 afferent ablation acutely four-to-six weeks after MI. Acute epidural RTX administration in chronically infarcted pigs was well tolerated, with no observed hemodynamic instability. Within 4 hours of administration, we observed improvements in vagal BRS, increased ventricular refractory period, attenuated pacing-induced dispersion of repolarization, and lower VT/VF inducibility. Taken together, these data suggest that, in the setting of chronic MI, cervicothoracic nociceptive spinal afferents, perhaps by increasing efferent outflow, are key contributors to the maintenance of pathological vagal dysfunction and cardiac electrical instability, thereby supporting their role in VT/VF susceptibility.

These findings carry important clinical implications. Given the selectivity of RTX for nociceptive TRPV1-expressing neurons, it is currently under investigation in clinical trials for chronic and refractory pain syndromes (NCT00804154; NCT02522611), highlighting its translational potential. In addition to showing that nociceptive cervicothoracic spinal afferents are critical to the pathogenesis and maintenance of autonomic remodeling in the setting of chronic MI, our data support the clinical potential of acute and chronic, selective, spinal afferent ablation after MI. Importantly, while epidural RTX was well tolerated hemodynamically in both healthy and infarcted animals, initial administration of RTX was associated with a transient sympatho-excitatory phase that peaked within 1 hour. In our cohort of animals where RTX was delivered prior to MI, this was accompanied by increased incidence of ventricular arrhythmias in the early MI convalescent period (∼2 hours post-RTX).

Overall, the present study supports selective spinal ablation of TRPV1-expressing nociceptive afferents as a strategy to improve sympatho-vagal imbalance and acutely reduce VT/VF inducibility in chronically infarcted hearts. These findings are in line with prior work demonstrating the acute anti-arrhythmic effects of direct RTX application on thoracic DRGs in a porcine model of acute ischemia,^29^ and extend these observations from the acute ischemic setting to the chronic development and maintenance of anti-arrhythmic benefit and mitigated autonomic remodeling in the setting of chronic MI. Future studies are warranted to further determine the durability of, and extent to which cervicothoracic epidural RTX after chronic MI can *reverse* or *modulate* pathological cardiac autonomic remodeling. As mentioned above, the reductions in VT/VF-inducibility in chronic RTX-treated animals were comparable to the anti-arrhythmic benefits reported with thoracic epidural anesthesia using bupivacaine^64^ and cardiac sympathetic denervation in patients with refractory VT/VF and electrical storm^119,120^. Advantages of epidural RTX over these therapies, would be its non-surgical approach, and the specificity of its target, as it leaves both mechanosensory afferents (in contrast to thoracic epidural anesthesia) as well as cardiomotor efferents intact.

Several limitations should be acknowledged. Although the initial sympathoexcitatory responses RTX and post-mortem histology supported successful local TRPV1 nociceptive afferent ablation, acute confirmation of ablation in vivo was not possible. In addition, given epidural administration at C7-T1 vertebral levels, effects of RTX in this study may represent an underestimate of its potential efficacy, as complete blockade of *all* cervicothoracic spinal afferent nerves was difficult to confirm. However, the blockade herein was sufficient to mitigate VT/VF inducibility and reduce post-MI autonomic remodeling.

While RTX was administered at the C7-T1 levels only, effects on other organ systems cannot be completely ruled out. Nevertheless, as renal^121–123^, splenic^124^, and hepatic^125–127^ spinal afferents originate from low thoracic/lumbar levels, and prior work indicates limited spread of epidurally-administered compounds^36–38^ (including epidural RTX), it is unlikely that these organs were affected significantly by RTX administered at the cervicothoracic level. Consistent with this, plasma aspartate transaminase (AST) and alanine aminotransferase (ALT), clinical indices of liver health, were similar between cRTX and vehicle-treated animals. Although pulmonary spinal afferents run through similar thoracic levels as cardiac spinal afferents, pulmonary innervation by spinal TRPV1-expressing nociceptive neurons is sparse^128^, and largely confined to large airways. Furthermore, activation of these pulmonary afferents has been shown to result in large airway smooth muscle relaxation,^129–135^ and therefore, unlikely to account for the cardiac electrophysiologic and autonomic results observed in this study. More distally, we did not observe effects on the T12-L1 spinal cord/DRG that innervate the kidneys, consistent with renal-independent benefits of RTX. Nevertheless, as assessment of CGRP innervation of pulmonary, splenic, renal or hepatic tissue was not performed in this study, potential off-target effects or benefit, which may have contributed to the results observed, cannot be completely ruled out. Finally, inflammatory-mediated cardiac remodeling was not directly characterized in this study. As such, the improved autonomic tone and electrical stability observed with RTX treatment may reflect a combination of direct neurally-mediated benefit and indirect anti-inflammatory benefit.

Lastly, in this study, CGRP was used as a surrogate for TRPV1-expressing nociceptive neurons/fibers, as direct TRPV1 immunolabeling was unreliable in our hands. However, activation of TRPV1 has been reliably shown to evoke the release of neurotransmitters such as CGRP^136–142^. Therefore, CGRP has been widely used as a marker for TRPV1-expressing afferent neurons.

In conclusion, cervicothoracic spinal/sympathetic nociceptive afferent signaling plays a significant role in the occurrence of the multitude of reported pathological autonomic changes after chronic MI. In chronically infarcted animals, ablation of C7/T1 spinal afferents prior to MI was associated with preserved vagal function, reduced glial activation and inflammatory signaling, and improved electrophysiological stability at four to six weeks post-MI. These changes coincided with lower dispersion of repolarization in border zone regions, areas known to provide both trigger and substrate for VT/VF, and ultimately with reduced VT/VF inducibility. The anti-arrhythmic benefits observed with epidural RTX in the chronic MI setting may therefore reflect a combination of dampened inflammatory/stress pathways together with preservation of parasympathetic tone and vagal reflex, limiting electrical heterogeneity.

Moreover, in a cohort of animals where RTX was administered instead 4-6 weeks following myocardial infarction, acute cervicothoracic epidural RTX was associated with improved vagal function and reduced VT/VF inducibility, suggesting potential utility for patients presenting with refractory VT/VF in the setting of a prior MI. Collectively, these findings offer important insights on the mechanisms underlying the chronic pathological autonomic remodeling and susceptibility to ventricular arrhythmias after MI.

## METHODS

### Ethical approval

Animal care was performed according to the National Institutes of Health Guide for the Care and Use of Laboratory Animals. All experimental protocols were approved by the UCLA Institutional Animal Care and Use Committee. All animals had an acclimatization period of seven days at our animal facility, prior to undergoing any experiments. The temperature was regulated between 16 and 18 °C. Swine were housed alone or in pairs, and boxes were enriched with chains, balls, and additional playing materials. Pigs were exposed to dark–light cycles of twelve hours with artificial lighting between 7 and 19 o’clock. Drinking water was supplied ad libitum and pigs were fed regular chow (LabDiet Mini-Pig; Item #55894).

In total, 57 male Yorkshire pigs (S&S Farms) were included in the study. Of these, three were healthy animals (Control; *n*=3) initially used to assess hemodynamic effects of acute epidural RTX. Of the 54 infarcted animals, 39 pigs survived the MI creation and the follow up four-to-six-week post-MI period to terminal studies. Nineteen animals were randomized to epidural RTX administration prior to MI (cRTX), of which 11 survived the acute MI and the four-to-six-week period post-MI, and 18 animals were randomized to vehicle (underwent thoracic epidural catheter placement without RTX administration) prior to MI, of which 13 survived the acute MI and the four-to-six-week period following MI. A fourth group of 17 animals underwent MI creation, of which 15 survived the acute MI and the four-to-six-week period post-MI, and were used for evaluation of the effects of acute cervicothoracic epidural RTX in chronically remodeled infarcted animals, Figure 1A. In cRTX- and vehicle-treated animals, terminal experiments were performed four to six weeks after epidural RTX injection and MI creation. Vehicle-treated animals had an epidural catheter placed and contrast/saline (vehicle) injected without RTX, 2-3 hours prior to induction of MI, with terminal assessments also performed at four-to-six weeks post-MI and saline injections (Figure 1).

On the other hand, in the aRTX cohort, chronic MI was induced, and animals were evaluated four to six weeks after. The epidural catheter was placed and RTX administered with the same approach. Hemodynamic and electrophysiological parameters were acutely assessed before and for the four hours following administration of epidural RTX.

Not all animals underwent the same interventions at the time of terminal studies due to the prolonged length of experiments and potential for micro- or macro-dislodgment of neural recording electrodes from the VIVGP with cardioversion of VT/VF or rinsing with saline after epicardial bradykinin/capsaicin application. Hence, animal numbers are indicated with each intervention under each section of the results.

Animals were monitored throughout the study by laboratory staff under veterinary oversight. Humane endpoints followed institutional Animal Research Committee policy, consistent with IACUC guidelines, and included chronic weight loss exceeding 20% of baseline body weight, a body condition score below 2, a Grimace scale score above 2, uncontrollable seizures, incoordination or paralysis, respiratory distress, decreased mobility interfering with normal eating or drinking, weak response to external stimuli, or uncontrolled bleeding; no animals met these criteria. Terminal experiments were open-chest procedures, and euthanasia was performed by induction of ventricular fibrillation with low-voltage direct current while animals were maintained in a deep surgical plane of anesthesia under inhaled isoflurane (4–5%), with death confirmed by loss of heartbeat over a minimum of 10 minutes together with standard clinical criteria. Animals reaching a humane endpoint prior to the terminal study would have been euthanized with intravenous pentobarbital or pentobarbital solution (≥100 mg/kg), with death confirmed by cessation of breathing, loss of pulse, and absence of corneal reflex.

### Creation of myocardial infarcts

MI was created percutaneously under fluoroscopic guidance, as previously described^49^. Briefly, animals (40.4±0.7 kg; 93±13 days of age) were sedated with tiletamine-zolazepam (4-8 mg/kg, intramuscular), intubated, and placed under general anesthesia (isoflurane 1-2%, inhaled). Pre-operative analgesic care consisted of buprenorphine (0.02 mg/kg; intramuscular) and carprofen (4.4 mg/kg; intramuscular). A coronary guide wire was introduced *via* the femoral artery into the left anterior descending coronary artery (LAD), and a balloon-tipped angioplasty catheter was advanced over an angioplasty wire past the first diagonal branch of the LAD and placed at the second diagonal branch. Next, the balloon was inflated and 3-4 ml polystyrene microspheres (Polybead, 90 μm, Polysciences, Warrington, PA) were injected through the angioplasty balloon lumen into the distal LAD. The balloon was deflated, and MI confirmed by lack of distal LAD flow on coronary angiography coupled with ST-segment changes (Figure 1B-C). Post-operative, animals were continuously monitored until they were well awake and eating, after which they were checked on every 12 hours for the first 3 days, and every day after that. Post-operative analgesia was continued for 72 hours post-experiment, and consisted of buprenorphine (0.02 mg/kg; intramuscular; twice a day) and carprofen (4.4mg/kg; intramuscular; once a day). Animals were then survived for four to six weeks.

### Cervicothoracic (C7-T1) epidural nociceptive denervation with resiniferatoxin

Pigs were placed in the lateral decubitus position. Using a paramedian approach, a 17-gauge Tuohy needle was inserted at the T5-T6 vertebral level, and the epidural space identified by standard loss-of-resistance under fluoroscopic guidance. Correct placement of the catheter in the epidural space was then confirmed by contrast injection under fluoroscopic guidance. A 19-gauge open-end epidural catheter (Teleflex Inc, Wayne, PA) was then advanced superiorly to the C7-T1 vertebral level (Figure 1F) and contrast again injected to confirm position. RTX or saline with contrast was injected into the epidural space. All cRTX animals (those receiving RTX prior to MI and studied four-to-six weeks later) received 1.2 µg/kg of RTX in 1 mL. In the chronic infarcted animals that received RTX four to six weeks after MI acutely, 7/15 received 1.2 µg/kg of RTX while 8/15 a lower dose of 0.6 µg/kg of RTX (also in 1 mL) to evaluate any differences in hemodynamic or electrophysiological responses. As no hemodynamic or electrophysiological differences were observed between the two groups (Supplemental Figure 9), data was pooled for analyses involving the aRTX animals.

### Animal preparation

For terminal studies, animals (49.8±0.1 kg; 127±10 days of age) were sedated with tiletamine-zolazepam (4-8 mg/kg, intramuscular) and intubated. General anesthesia was induced with isoflurane (1-2%, inhaled) and transitioned to α-chloralose (50 mg/kg initial bolus, then 20-30 mg/kg/hr. infusion) prior to autonomic or electrophysiological testing. Sheaths were placed in bilateral femoral veins and arteries for saline infusion, drug administration, pressure monitoring, and introduction of electrophysiological catheters.

### Intracardiac echocardiography

Prior to any intervention, an intracardiac echocardiography (ICE) probe (ViewFlex Xtra ICE Catheter, Abbott, Minneapolis, USA), connected to the ViewMate Multi Ultrasound System (Abbott, Minneapolis, USA), was introduced *via* the right femoral vein into the right atrium in 10 vehicle- and 10 RTX-treated (cRTX) animals. Upon placement in the right atrium, the probe was slowly retracted and rotated to obtain a view similar to the parasternal short axis view. Recordings of the LV were obtained at the level of the papillary muscles. Images were analyzed offline using Sante DICOM Viewer Lite (Santesoft Ltd, Nicosia, Cyprus) to obtain diastolic and systolic LV area and wall thickening measurements.

### Ventricular hemodynamic measurements

A 5-Fr Millar pressure-conductance catheter was introduced *via* the femoral artery and placed in the LV for continuous pressure. Raw signals were digitized by a CED Power1401 and analyzed using Spike2 (Cambridge Electronic Design, v10.08). A continuous 12 lead ECG was obtained via a CardioLab Recording System (GE Healthcare).

### Ventricular electrophysiological measurements

All animals underwent median sternotomy to expose the heart. A 56-electrode sock, connected to a GE CardioLab system, was placed over the ventricles for continuous local unipolar epicardial electrogram recordings (band pass filtered 0.05-500 Hz, Figure 1D). Activation time (AT) and repolarization time (RT) were measured by customized software (iScaldyn; University of Utah) from these unipolar electrograms as the intervals from onset of ventricular activation to the minimal dV/dt of the depolarization wave-front or maximal dV/dt of the repolarization wavefront, respectively (Figure 1E). Activation recovery interval (ARI), a surrogate for action potential duration,^143^ was calculated as the difference between RT and AT, and corrected for differences in HR using the Bazett formula. Global dispersion in RT (DoR) was calculated as the RT variance across all sock electrodes, whereas regional DoR was calculated as the variance across electrodes assigned to regions based on bipolar voltage mapping (below).

Bipolar voltage mapping was performed in epidural cRTX (*n*=11) and vehicle (*n*=13) animals using a standard 2-2-2 duodecapolar catheter (Abbott, Minneapolis, USA) to delineate scar, border zone and viable regions. For this purpose, the duodecapolar catheter was advanced between the epicardium and the sock electrode and the bipolar voltage underneath each respective electrode was measured. Using standard voltage criteria regions were defined as either scar (0.05 mV<voltage<0.5 mV), border zone (0.5 mV<voltage<1.5 mV), or viable (voltage>1.5 mV) myocardium^61^.

### Evaluation of cardiac autonomic function

#### Baroreflex sensitivity

Vagal baroreflex sensitivity was tested in vehicle-treated animals (epidural with saline/contrast only followed by MI; *n*=13), cRTX (epidural with RTX followed by MI with evaluation four-to-six weeks post-MI, *n*=11), aRTX (MI followed by epidural RTX four to six weeks later; *n*=12) by bolus injection of phenylephrine (3-5 μg/kg, IV) to evoke a 30-40 mmHg increase in systolic pressure. Vagal BRS was measured hourly for four hours in aRTX animals, while it was measured once in control, vehicle and cRTX animals. The slope of the linear regression describing the beat-to-beat relationship between RR-interval and LV systolic pressure (LVSP) was used to quantify baroreflex sensitivity (BRS)^144^.

#### Cardiac nociceptive stimulation

Epicardial application of bradykinin (1.06 mg/mL) and capsaicin (0.03 mg/mL) in vehicle (*n*=10) and cRTX (*n*=11) animals was used to characterize autonomic responses to epicardial stimulation of chemosensitive, nociceptive afferents. Chemicals were individually applied (15 mL over 10 seconds) and thoroughly washed off with 500 mL of warm saline 1-minute after start of application. A 15-30 min wait period was allowed for hemodynamic parameters to return to baseline prior to subsequent interventions.

#### Extracellular neural recording from the intrinsic cardiac nervous system

Custom-made 16-channel linear microelectrode arrays (MicroProbes for Life Science; 25-µm-diameter platinum/iridium electrodes, 16 electrodes/probe, 375-µm interelectrode spacing) were used for *in vivo* extracellular neural recordings of the VIVGP (Figure 4), as previously described^48,49,66^ in 5 vehicle and 6 epidural cRTX animals. In short, the probe was gently advanced into the epicardial fat pad and serially connected to a head-stage preamplifier and a 16-channel preamplifier (Model 3600, A-M Systems, Sequim, WA). All signals were continuously recorded and digitized (Cambridge Electronic Design) at a sampling frequency of 20 kHz, and band-pass filtered (0.3 - 3 kHz). Offline processing and analyses of neural signals was performed using Spike2 software (Cambridge Electronic Design, v10.08), as previously described^48,49,66^. Artifacts (recognized as simultaneous waveforms on all neural recording channels) were removed, and neuronal spikes were identified using a threshold of 2 times signal-to-noise ratio. Spike sorting was performed using principal components, cluster on measurements, and K-means clustering analysis to identify unique neuronal waveforms^47,145^. Efferent postganglionic parasympathetic neurons in the VIVGP were identified based on their responses to left or right-sided VNS. Briefly, bipolar spiral cuff electrodes (LivaNova, PLC) were placed around each cervical vagus, and the VIVGP activation threshold current, defined as the VNS current needed to evoke a 10% decrease in heart rate (20 Hz, 1 ms), was determined. Each cervical vagus was then stimulated separately for 1 minute at 1 Hz (1 ms, 1× VIVGP activation threshold current). At least 20 minutes of recovery time was allowed between the two stimulations. Baseline activity during the 2 minutes before left or right VNS was compared with the 2 minutes at the start of stimulation (1 minute during VNS and 1 minute after) using the Skellam statistical test^146^. Neurons that showed significant changes in firing activity (as compared by the Skellam test) upon left or right VNS were identified as postganglionic parasympathetic neurons, while neurons that did not show significant alterations in their activity were considered to be non-VNS responsive. Basal activity of both VNS-responsive and non-VNS responsive (1 minute) was compared between vehicle (*n*=5) and epidural cRTX (*n*=6) animals.

#### Effective refractory period measurements and VT/VF inducibility

Atrial and ventricular effective refractory periods (ERP) were measured by extra-stimulus pacing at a drive cycle length (CL) of 450 msec, with S2 decremented by 5 msec, using a pacing catheter placed on the epicardial left atrial appendage and in the right ventricular (RV) apex from the right femoral vein.

VT inducibility was tested in epidural cRTX (*n*=11), vehicle (*n*=13), epidural aRTX (*n*=12) animals. Inducibility was assessed by programmed stimulation (as is standard in electrophysiology laboratories for testing of inducibility in patients with heart disease)^147^ by an 8-beat drive train (at CL of 450 ms) followed by an S2 extra-stimulus, which was decremented by 10 msec down to a CL of 200 msec or ERP, whichever occurred first. If no VT/VF was induced, a CL of 20 msec above ERP was selected for the extra-stimulus to ensure ventricular capture and the next extra-stimulus (up to S4) was added. VT/VF inducibility was defined as the occurrence of sustained VT (>30 seconds) or VF requiring defibrillation. Inducible animals were cardioverted if VT/VF did not terminate after 30 seconds. VT inducibility was tested from RV endocardium and, if non-inducible, also from the LV anterior epicardial border zone region. For acute RTX animals, the same site that induced VT/VF pre-RTX administration was used post-RTX administration to induce VT. Ventricular pacing threshold were checked and maintained pre- *vs* post-RTX administration (Supplemental Figure 8).

### Histopathological assessment

LV tissue, macroscopically and electrically (by bipolar voltage criteria, defined as voltage > 1.5 mV^61^) classified as viable, as well as C7-T1 and T12-L1 spinal cords and DRGs were collected from cRTX (*n*=5-7) and vehicle-treated infarcted animals (*n*=5), fixed in 4% paraformaldehyde, and embedded in paraffin. Tissue was sectioned at 5 µm, deparaffinized, rehydrated, and epitopes unmasked at 90 °C in EDTA buffer (Abcam, ab64216). Slides were blocked and incubated overnight at 4 °C with goat anti-calcitonin gene-related peptide (CGRP; 1:1000; Abcam, ab36001), mouse anti-glial fibrillary protein (GFAP; 1:1000; Invitrogen, ASTRO6), rabbit anti-ionized calcium-binding adaptor molecule 1 (Iba1; 1:1000; Biocare Medical, CP 290 A, B), sheep anti-tyrosine hydroxylase (TH; 1:500; MilliporeSigma, AB1542), rabbit anti-purine P2X3 receptor (1:200; Neuromics, RA10109), rabbit anti-Piezo-type mechanosensitive ion channel component 2 (PIEZO2; 1:400; ThermoFisher, PA5-72975) and/or rat anti-cluster of differentiation 3 (CD3; 1:1000; Abcam, ab11089). CGRP was used as a surrogate for TRPV1 expression as it is released by nociceptive sensory/afferent neurons upon TRPV1 activation^138^. Sections were incubated for 2 hours at room temperature with Alexa Fluor 488–donkey anti-goat IgG (1:200; Invitrogen, A-11055), Alexa Fluor 555–donkey anti-mouse IgG (1:400; Invitrogen, A-31570), Alexa Fluor 488–donkey anti-rabbit IgG (1:400; Invitrogen, A21206), Alexa Fluor 647–donkey anti-sheep IgG (1:400; Invitrogen, A-21448), and/or CF405S–donkey anti-rat IgG (1:400; Biotium, 20419), respectively and mounted with Antifade Mounting Medium (Vector Laboratories, H-1000-10). Slides were imaged on a Zeiss LSM880 at 20× and 63× magnifications and processed with Zen 2 software (Zeiss). In the spinal cord, CGRP, GFAP, IBA1, and CD3 immunoreactivity were quantified as the fractional area of the dorsal horn. In the DRG, GFAP, CD3, and IBA1 were quantified as the fractional area and CGRP quantified as percentage of neurons in the section containing the greatest area of neurons, respectively. All image analysis was performed with ImageJ (NIH)

### Measurement of plasma liver enzymes and inflammatory markers

For evaluation of liver function, femoral artery blood was collected at the time of terminal studies (4-6 weeks after vehicle or RTX injection and MI) to measure plasma interleukin-12 (IL-12), tumor necrosis factor α (TNF-α), aspartate aminotransferase (AST) and alanine aminotransferase (ALT) concentrations in animals treated with epidural vehicle or RTX prior to MI. Blood was collected into K_2_ EDTA blood collection tubes (BD Vacutainer), and immediately centrifugated at 1500 *g* for 15 minutes. Plasma was separated, snap-frozen in liquid nitrogen and stored at −80°C until assay. IL-12 (P1240, sensitivity 18.2 pg/mL, Quantikine), TNF-α (PTA00, sensitivity 5pg/mL, Quantikine), AST (MBS739883, sensitivity 0.1ng/mL, MyBioSource) and ALT (MBS266004, sensitivity 0.06ng/mL, MyBioSource) levels were measured by ELISA, according to manufacturer’s instructions.

### Plasma proteomics

#### Sample preparation

After access to the femoral artery was obtained and prior to performing any autonomic or electrophysiological testing, baseline blood was collected in EDTA blood tubes and immediately placed on ice. Samples were centrifuged and plasma was separated and stored in −80 °C. Plasma samples (n=9 for Vehicle + MI; n=10 for cRTX + MI) were prepared by the University of California, Los Angeles (UCLA) Proteome Research Center, using the Mag-Net^148^. Dried peptides were resuspended in 5% formic acid and separated using PepSep C18 reverse-phase column (150□mm × 150□µm, 1.7□µm particle size) heated to 59□°C. Fractionation was conducted on a Thermofisher Vanquish Neo UHPLC system using a trap-and-elute workflow and a 15 minute gradient of increasing acetonitrile (5% acetonitrile to 80% acetonitrile) delivered at a flow rate of 2.45□µL/min. MS/MS spectra were collected using a data-independent analysis (DIA) acquisition method on ThermoFisher Orbitrap Astral mass spectrometer^149,150^. MS/MS spectra were acquired using sequential 4 m/z windows tiled over a m/z range of 380–980 at 80,000 resolution with a normalized HCD energy of 25%, a normalized AGC target of 500%, and a 7□ms maximum injection time. Data were analyzed using the DIA-NN algorithm to search an in silico spectral library generated from the Sus scrofa UniProt reference proteome (ID: UP000008227). Peptide and protein identifications were filtered using an estimated false discovery rate of less than 1%^151^. Comparison testing between conditions was performed using Fragpipe-Analyst platform using DIA-NN-generated label-free protein abundances^152^.

#### Data analysis

Data analysis was performed in R. Raw counts were normalized with random-forest normalization using R. Differentially expressed proteins were identified using a two-sided Student’s t-test. Proteins with a *p-value* < 0.05 were considered statistically significant, and those with *|log2(fold change)|* > 0.5 were identified as differentially expressed proteins. Pathway analysis was performed against the Gene Ontology (GO) database using Rapid Integration of Term Annotation and Network (RITAN, v3.20)^153^. A two-sided *p*-value < 0.05 was used to determine statistically overrepresented pathways.

### Analysis of plasma inflammatory cytokines

Femoral artery blood was collected at the time of terminal experiment (4-6 weeks after vehicle or RTX injection and MI) to measure plasma tumor necrosis factor-α (TNF-α), interleukin-1β (IL-1β), interleukin-6 (IL-6), interleukin-10 (IL-10), and interleukin-12 (IL-12). Blood was collected into K2 EDTA blood collection tubes (BD Vacutainer), and immediately centrifugated at 1500 g for 15 minutes. Plasma was separated, snap-frozen in liquid nitrogen and stored at −80°C until assay. Circulating TNF-α, IL-1β, IL-6, IL-10, and IL-12 were assessed via Porcine Luminex Discovery Assay (R&D Systems) according to the manufacturer’s instructions.

### Statistical Analyses

Data are presented as mean ± SEM. For cRTX versus vehicle comparisons, each data point represents an independent biological replicate (one animal); for acute RTX (aRTX) experiments, parameters were measured repeatedly within the same animal before and after RTX administration, and analyzed using paired/repeated-measures tests. After confirmation of normality, paired two-tailed Student’s *t*-test or Wilcoxon signed rank test was used to compare pre- and post-RTX parameters in epidural aRTX animals, depending on Gaussian distribution. Immunohistochemical data were analyzed using unpaired Student’s t-tests or Mann-Whitney U test (depending on Gaussian distribution) with df = n_1_+n_2_−2 for each comparison (derived from the group sizes stated per panel). Unpaired analysis of variance (ANOVA) was used for intergroup comparisons (vehicle *vs* epidural cRTX *vs* Control) and changes in hemodynamic parameters over time were compared using repeated measures ANOVA. Electrophysiological parameters were compared using unpaired Student’s t-tests or Mann-Whitney U test (depending on Gaussian distribution) for epidural cRTX *vs* vehicle animals or repeated measures ANOVA in case of serial comparisons in acute RTX studies. BRS data was only included for analysis if R^2^ was greater than 0.8 for the slope of the regression line relating blood pressure to heart rate and compared using the Mann-Whitney U test (for group comparisons) or the Friedman test (for serial comparison in acute RTX experiments). Comparison of VT/VF inducibility was performed using the binomial exact test. *P-value* < 0.05 was considered statistically significant.

## DATA AVAILABILITY

The mass spectrometry proteomics data have been deposited to the ProteomeXchange Consortium via the PRIDE partner repository under accession code MV000099727 (https://massive.ucsd.edu/ProteoSAFe/dataset.jsp?task=9d63769e90804f8091546cfca56052cb). Source data underlying all figures, including individual data points, are provided with this paper in the Source Data file and Supplementary data 1 and 2. Raw acquisition files (electrophysiological recordings, neural recordings, and confocal microscopy images) are large and are available from the corresponding author on request, owing to file-size and format constraints; their summary measurements are fully reported in the Source Data file.

## CODE AVAILABILITY

Custom code used to process the extracellular neural recordings and to analyze the plasma proteomic data (differential expression, and pathway enrichment) is available at GitHub (https://github.com/jdthoang/vanWeperenV_2026_EpiduralRTX/) and archived at Zenodo (https://doi.org/10.5281/zenodo.20711756). Pathway analysis was performed with RITAN (v3.20) in R (v4.4.2). Customized software, iScaldyn (University of Utah), was used to compute activation-recovery intervals and is freely available upon request from Robert L. Lux. Image quantification was performed in ImageJ (NIH, v1.54). No other custom code was generated in this study.

## Abbreviations

ARI: activation-recovery interval
AT: activation time
BRS: baroreflex sensitivity
CD3: cluster of differentiation 3
CGRP: calcitonin gene-related peptide
CL: cycle length
DoR: dispersion of repolarization time
ECG: electrocardiogram
ERP: effective refractory period
GFAP: glial fibrillary acidic protein
HR: heart rate
LAD: left anterior descending
LV: left ventricle
LVSP: left ventricular systolic pressure
MI: myocardial infarction
RT: recovery time
RTX: resiniferatoxin
SCD: sudden cardiac death
TRPV1: transient receptor potential vanilloid 1
VIVGP: ventral interventricular ganglionated plexus
VNS: vagal nerve stimulation

## ACKNOWLEDGEMENTS

The authors are grateful for the help of the University of California, Los Angeles (UCLA) Proteome Research Center and Ms. Sakshi Fnu for her assistance with histological analyses.

## AUTHOR CONTRIBUTIONS

VvW, JDH, NJ, SA, CAC, KC, ZAL, ME, KA, and MV performed animal experiments. VvW, JDH, NJ, SA, KC, KA performed immunohistochemistry and slides were then imaged by VvW, JDH, and KA. VvW and JDH analyzed proteomics data. VvW, JDH, and MV drafted the manuscript. VvW, JDH and MV contributed to experimental design and data interpretation for aspects of data collected. Hemodynamic and electrophysiological analyses were performed by VvW, JDH, NJ, SA, CAC, KC, ZAL and ME. VvW, JDH and MV drafted the manuscript. All authors approved the final version of the paper.

## COMPETING RISK INTEREST STATEMENT

MV declares the following potential competing risks interest: She has shares in the following health-related companies: NeuCures Inc., Nference Inc., and Anumana Inc. She has a patent held of Regents of University of California - U.S Patent #20230330417A1: Neural modulation of autonomic nervous system to alter memory and plasticity of the autonomic network. Inventors are: Jeffrey L. Ardell, Marmar Vaseghi, Kalyanam Shivkumar. Patented. No aspect of the manuscript is covered by this patent.

All other authors have no competing risk interests to declare.

