## Supplemental figures for "Spinal nociceptive denervation impedes subsequent chronic autonomic remodeling after myocardial infarction in male swine"

**Supplemental Figure 1**

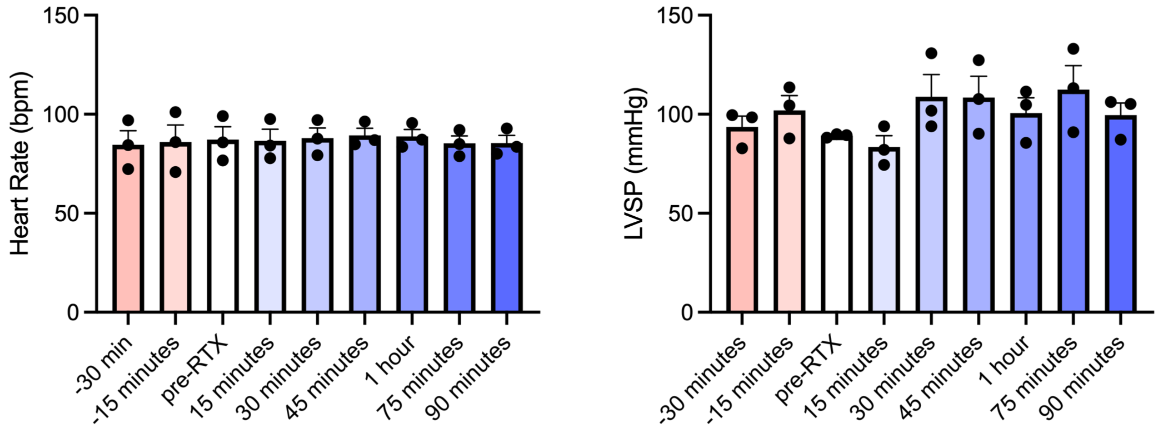


**Supplemental Figure 1. Temporal hemodynamic responses to epidural RTX administration in healthy control animals.** Absolute change in heart rate (HR) and left ventricular systolic pressure (LVSP) over time are shown for 3 healthy pigs. HR and LVSP, parameters which can be readily measured without the need for LV catheters, were assessed to determine the time to reach a steady state. A steady state was defined as no further significant changes in HR and LVSP over a thirty-minute period post-RTX administration. A steady state was reached approximately one hour after RTX administration.

**Supplemental Figure 2**

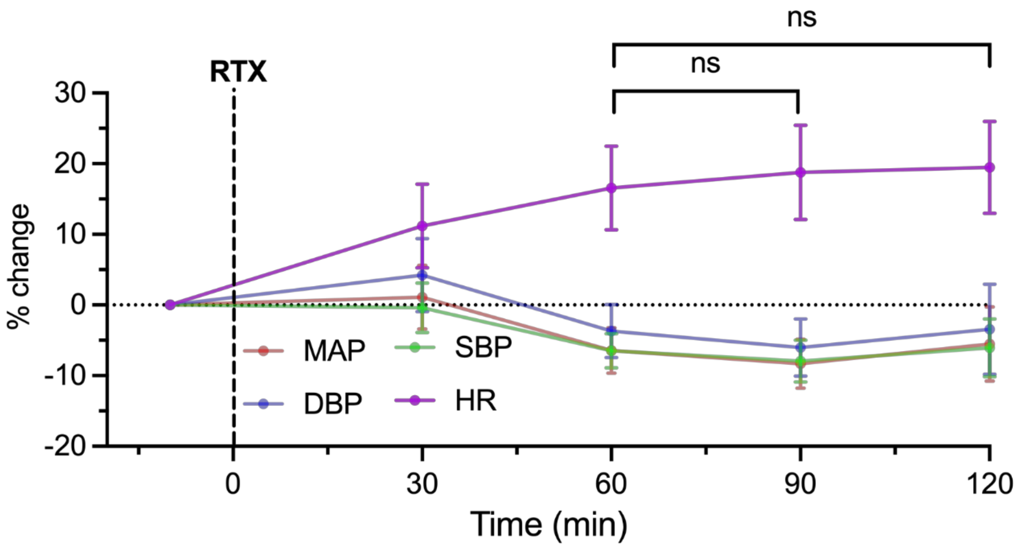


**Supplemental Figure 2. Temporal hemodynamic responses to acute epidural RTX prior to creation of myocardial infarction (MI).** Percentage change in HR, systolic (SBP) and diastolic blood pressure (DBP) and mean arterial pressure (MAP) over time are shown in the 11 animals that received epidural RTX 2-3 hours prior to creation of MI. At 60 minutes, hemodynamic stability was reached, as defined by no further significant changes in hemodynamic parameters over time.

**Supplemental Figure 3**

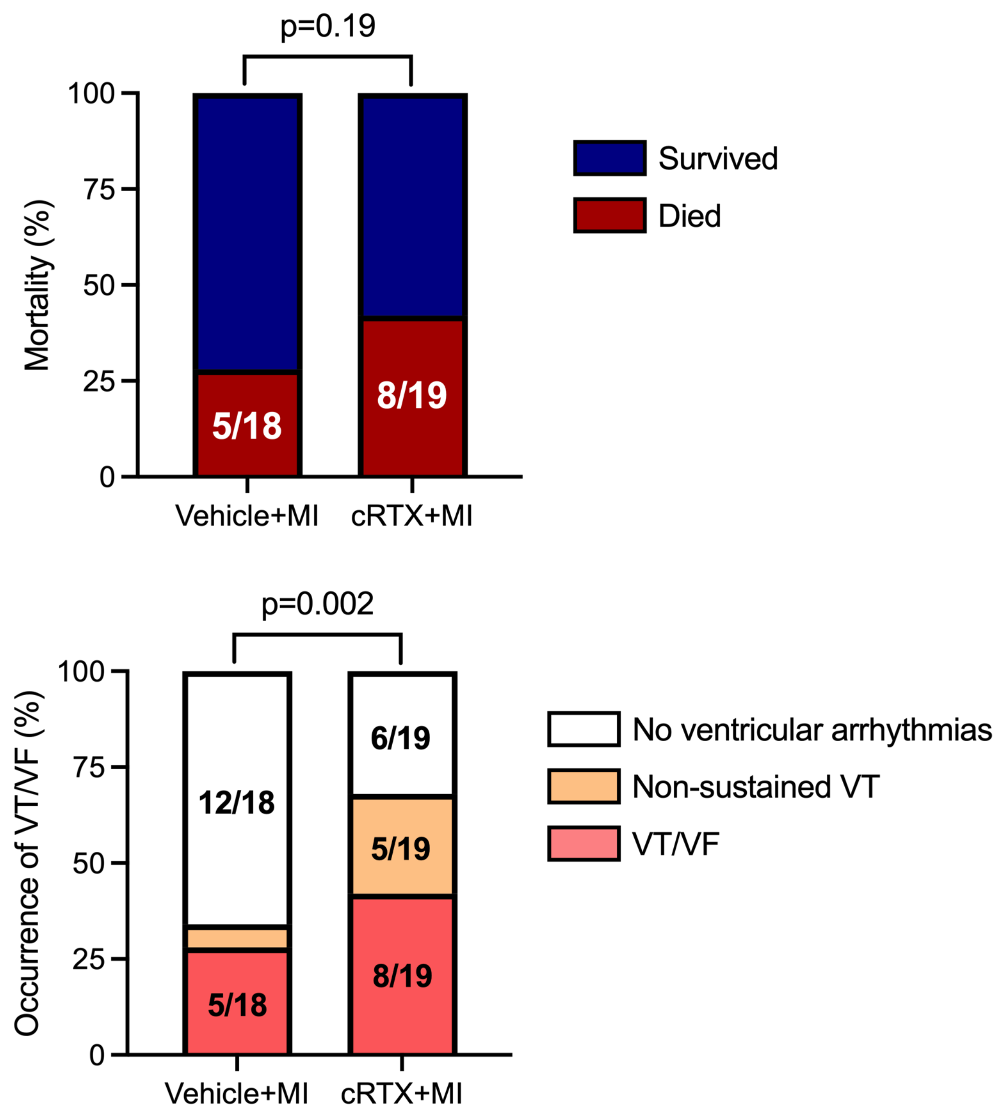


**Supplemental Figure 3. Mortality during creation of myocardial infarction (MI) in vehicle and cRTX animals.** There was no difference in mortality at the time of MI in animals that received epidural saline (Vehicle+MI) compared to those that received epidural RTX (cRTX+MI) prior to creation of myocardial infarcts. However, a greater number of animals in the cRTX group had spontaneous non-sustained VT or VT/VF at the time of MI creation. Comparisons were performed using the binomial exact test.

**Supplemental Figure 4**

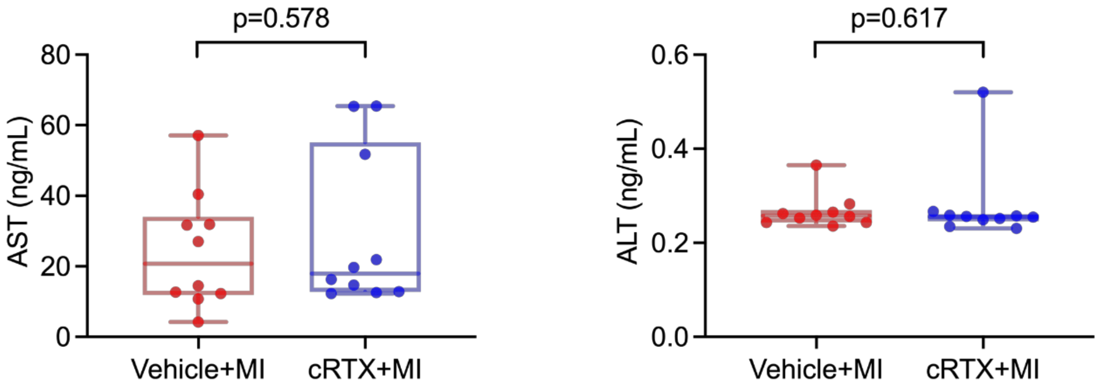


**Supplemental Figure 4. Assessment of liver function in vehicle *vs.* cRTX animals.** Comparison of the plasma levels of hepatic enzymes aspartate transaminase (AST) and alanine transaminase (ALT), which are clinically used to assess liver function, showed no significant differences in cRTX vs. vehicle-treated animals. Comparisons performed using the Mann Whitney test, $n$=10 animals per group.

**Supplemental Figure 5**

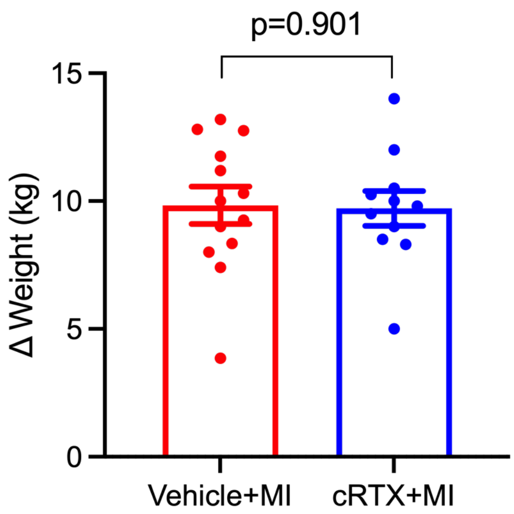


**Supplemental Figure 5. Assessment of weight changes in vehicle *vs.* cRTX animals.** The weight of animals in the cRTX (*n*=11) and vehicle-treated (*n*=13) groups was not different over the 4-6 weeks following myocardial infarction. Comparison performed using the unpaired Student's t-test.

**Supplemental Figure 6**

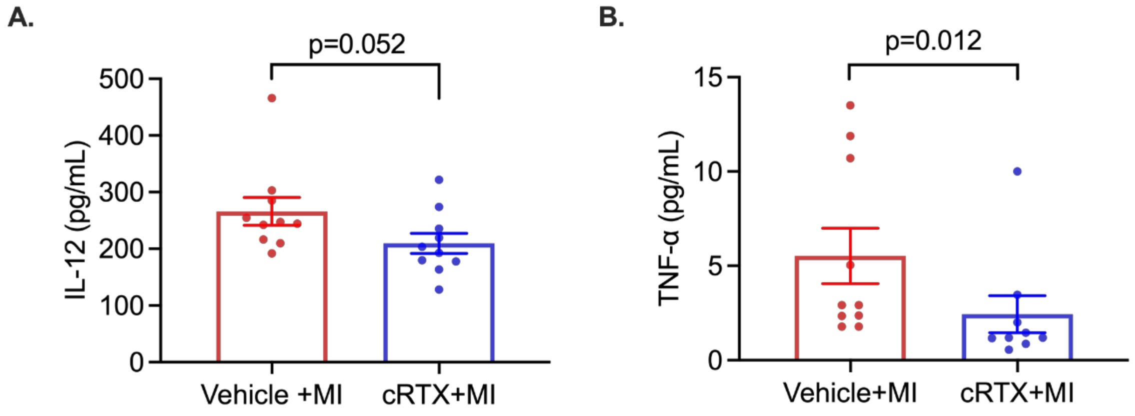


**Supplemental Figure 6. Assessment of plasma levels of interleukin-12 (IL-12) and tumor necrosis factor α (TNF-α) in vehicle *vs* cRTX animals. (A)** IL-12 levels were not statistically different between vehicle (*n*=10) *vs.* cRTX (*n*=10) animals. **(B)** TNF-α levels on the other hand, were significantly lower in cRTX (*n*=9) compared to vehicle-treated (*n*=10) animals. Differences between groups were assessed using the Mann Whitney test.

**Supplemental Figure 7**

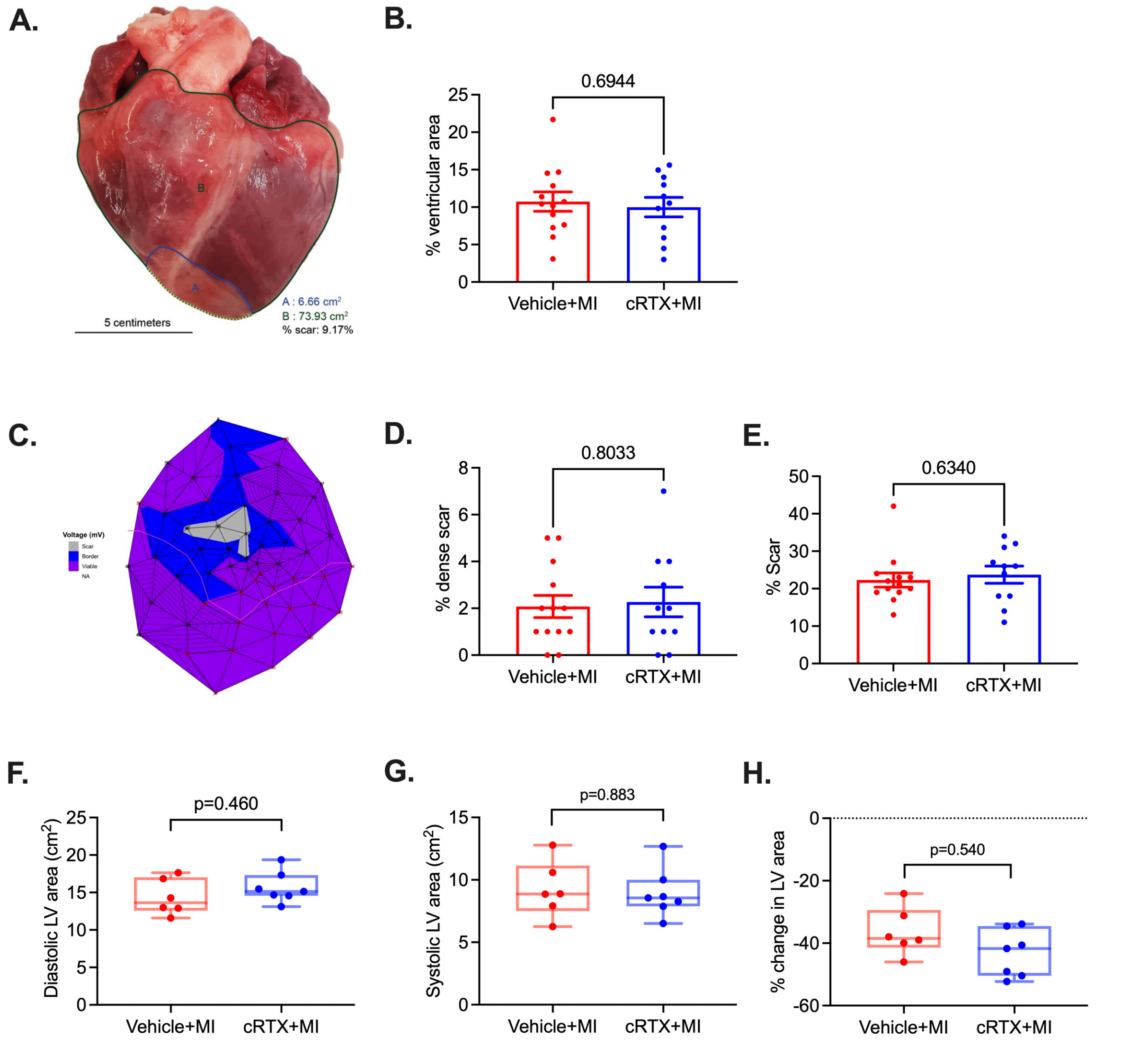


**Supplemental Figure 7. Assessment of myocardial infarct size. (A)** Post-mortem macroscopic assessment of scar size through ImageJ demonstrated **(B)** similar infarct sizes in RTX-treated (epidural cRTX+MI *n*=11) compared to vehicle-treated (*n*=13) animals. **(C)** Electrophysiological assessment of infarct size through epicardial bipolar voltage mapping demonstrated **(D)** similar dense scar sizes (defined as voltage < 0.5 mV) and **(E)** infarct size (infarcted area was defined as epicardial voltage < 1.5 mV to include border zone regions) between cRTX (*n*=11) and vehicle-treated (*n*=13), infarcted animals. Also functionally, assessed *via* echocardiography, there were no differences in **(F)** diastolic left ventricular (LV) area, **(G)** systolic LV area, and **(H)** percentage change in LV area (ejection fraction) in cRTX (*n*=10) versus vehicle-treated (*n*=10) infarcted animals. Comparisons performed using the unpaired Student's t-test.

**Supplemental Figure 8**

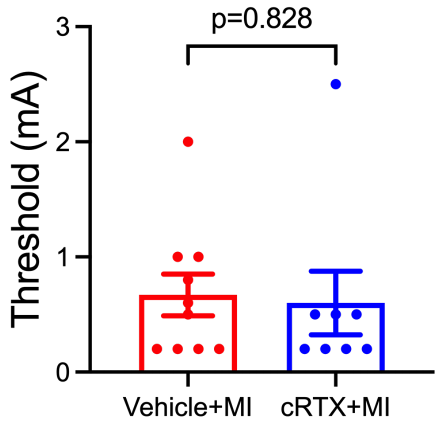


**Supplemental Figure 8. Comparison of ventricular pacing between vehicle and cRTX animals.** Right ventricular endocardial pacing thresholds were similar between vehicle-treated (*n*=10) and cRTX (*n*=8) animals. Comparison performed using the unpaired Student's t-test.

**Supplemental Figure 9**

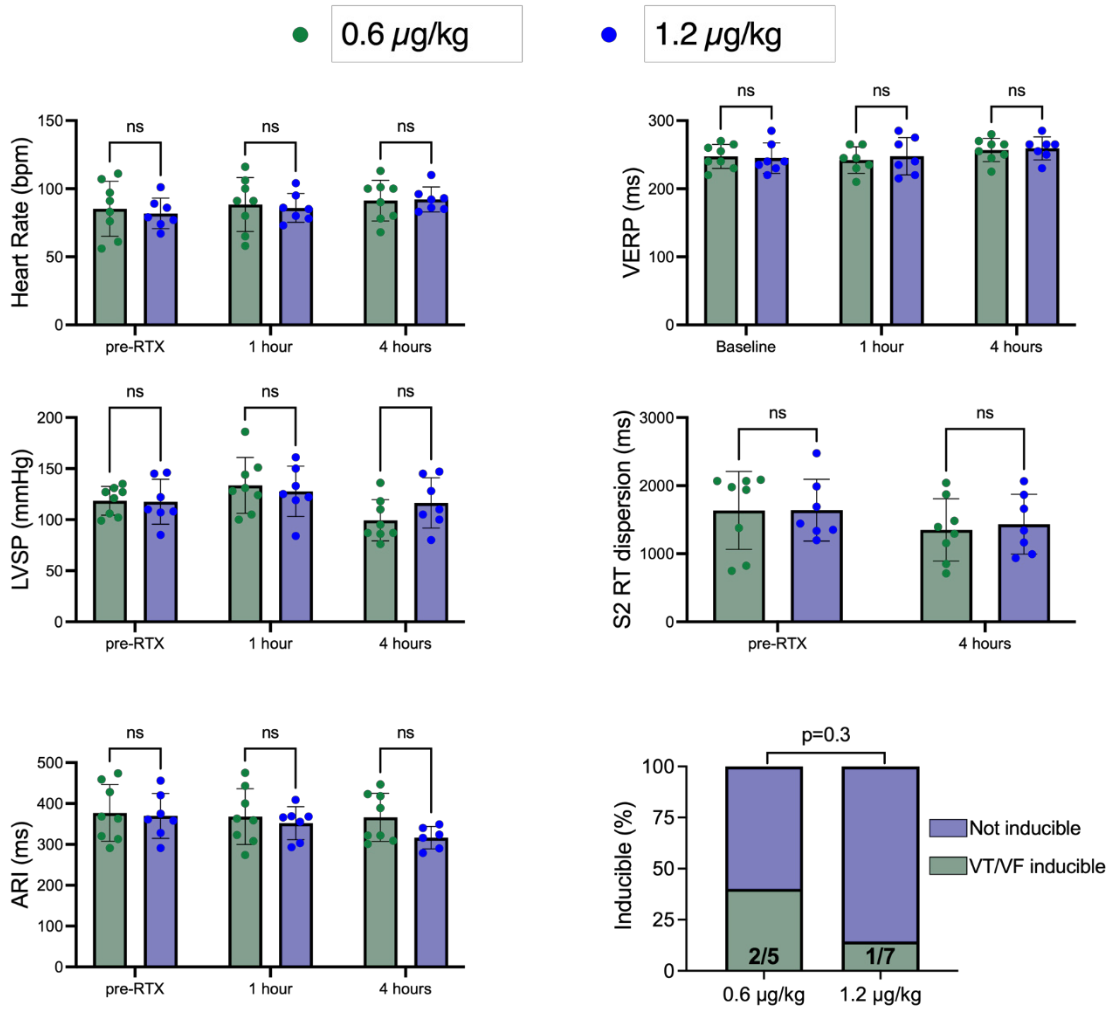


**Supplemental Figure 9. Assessment of hemodynamic and electrophysiological differences between animals receiving a single (0.6 μg/kg) or double (1.2 μg/kg) dose of epidural RTX in the setting of chronic myocardial infarction.** No hemodynamic or electrophysiological differences were observed between animals receiving a single (0.6 μg/kg; n=8) or double (1.2 μg/kg; n=7) dose of epidural RTX. All hemodynamic and electrophysiological comparisons were performed using the unpaired Student's t-test, VT inducibility was compared by the exact binomial test.
